# Contrasting Probabilistic and Intentional Accounts of Confidence in Perceptual Decisions

**DOI:** 10.64898/2026.03.24.714055

**Authors:** Ariel Zylberberg

## Abstract

The ability to evaluate one’s own knowledge states is often studied using paradigms in which participants make a decision and subsequently report their confidence. This structure has motivated hierarchical models in which confidence arises from a metacognitive process, distinct from the decision process itself, that estimates the probability that the choice is correct (Meyniel et al., 2015; Pouget et al., 2016; Fleming and Daw, 2017). Here, we contrast this framework with an alternative based on an intentional architecture (Shadlen et al., 2008). In this account, choice and confidence are determined simultaneously through a multidimensional drift–diffusion process, where each dimension represents one choice–confidence combination (Ratcliff and Starns, 2009, 2013). Choice, response time, and confidence jointly emerge when one of these accumulators reaches a decision bound. To adjudicate between these accounts, we fit both models to behavioral data from two perceptual tasks: a random-dots motion discrimination task with incentivized confidence reports, and a luminance discrimination task without feedback or incentives. The integrated model provided a superior fit for the incentivized motion task, whereas the hierarchical model more accurately captured behavior in the un-incentivized luminance task. These results suggest that confidence does not rely on a single computational mechanism, but rather its implementation may adapt to the specific demands and structure of the task.

## Introduction

The ability to evaluate one’s own knowledge states is typically studied using paradigms in which participants first make a decision and then report their confidence in its accuracy. Because confidence is defined relative to a specific choice—whether actual, hypothetical, or counterfactual—there is an inherently hierarchical dependency between the two variables. This logical dependency is often interpreted as reflecting a computational one: confidence is assumed to arise from a metacognitive process that reads out internal decision variables to estimate the probability that the choice is correct.

Most models that aim to explain choice, decision response time, and confidence fall within this hierarchical framework. In the Balance of Evidence (BoE) model (Vickers, 1979), for example, a decision emerges from a competition between evidence accumulators until one reaches a threshold; confidence is determined by a separate process that reads out the difference between the winning and losing accumulators at the time of response. Other models propose that confidence depends on response time (Audley, 1960; Zylberberg et al., 2012), a combination of response time and the BoE (Kiani et al., 2014; van Den Berg et al., 2016), the activity of choice-selective neural subpopulations (Zylberberg and Shadlen, 2025), the degree of consensus within those populations (Paz et al., 2016), the dispersion of the decision variable (Yeung and Summerfield, 2012), the attentional state (Brus et al., 2021), or an estimate of stimulus visibility (Hellmann et al., 2023). Despite their mechanistic differences, these approaches share a core architectural commitment: confidence is computed by a distinct process that accesses decision-related variables after a choice has been made.

These models were primarily developed for tasks in which confidence is reported sequentially, after the choice. Simultaneous-report paradigms—where choice and confidence are communicated through a single motor act—pose an additional challenge. In such cases, the total response time must reflect both the time required to reach the primary decision and the time needed for the metacognitive system to construct the confidence judgment from the corresponding readout. However, hierarchical models rarely specify the time required for this metacognitive process. In practice, when fitting these models to empirical data, the readout mechanism is often assumed to be instantaneous, such that the entire response time is attributed to the primary decision process (e.g., Kiani et al., 2014; van Den Berg et al., 2016; Vivar-Lazo and Fetsch, 2025).

Additional concerns about the validity of hierarchical models arise from studies comparing performance when confidence must be reported on a trial-by-trial basis versus when it is not required. Prompting for confidence can slow response times (Petrusic and Baranski, 2003; Dou et al., 2024), alter choice accuracy (Litwin et al., 2025; Double and Birney, 2024), and produce more polarized beliefs (Double et al., 2026). These findings suggest that confidence reports are not merely passive readouts of the decision state, but may instead form an integral component of the decision process itself.

An alternative to the hierarchical framework for simultaneous-report tasks emerges from the neurobiology of decision-making. Studies of simple perceptual decisions in non-human primates and rodents suggest that the brain does not resolve decisions by computing probability distributions over relevant variables. Instead, it represents and updates scalar decision variables, one for each alternative (Roitman and Shadlen, 2002; Steinemann et al., 2022; Hanks et al., 2015; DePasquale et al., 2021). These variables are encoded within an intentional reference frame: when the motor actions used to communicate the decision are known in advance, brain areas representing the intention to execute those actions reflect the accumulation of evidence during deliberation (Shadlen et al., 2008). The decision terminates when the accumulated evidence favoring one alternative exceeds a criterion (Ratcliff and McKoon, 2008; Gold and Shadlen, 2007). A natural extension of this intentional framework to simultaneous reports is to assume that evidence accumulates for each choice–confidence combination. In this view, confidence is not a second-order judgment but an intrinsic dimension of the same accumulation process that determines the choice.

Confidence models most consistent with this intentional perspective were proposed by Ratcliff and Starns (2009, 2013). These authors introduced race models in which each combination of choice and confidence level (e.g., “Left–High Confidence” or “Right–Low Confidence”) is implemented as a separate drift–diffusion process. The drift rate for each alternative is obtained by partitioning the evidence space that guides the decision. The process terminates when one of the drift-diffusion processes reach an upper criterion or bound, simultaneously resolving choice, response time and confidence. In this formulation, the distinction between first- and second-order judgments largely disappears, as choice and confidence emerge jointly from a single evidence-accumulation process.

In the present study, we empirically contrast these two conceptions of confidence using two perceptual tasks (see Chen et al. 2026 for a related comparison). The first is a random-dot motion discrimination task in which confidence was incentivized through a point system. The second is a luminance discrimination task with no feedback or explicit incentives.

## Results

We have structured the results into four sections. First, we briefly describe the experimental tasks. Second, we characterize the main behavioral patterns observed in each task. Third, we introduce the competing formal models proposed to explain these data. Finally, we evaluate the models’ performance by fitting them to the behavioral data.

### Tasks Overview

#### Random Dots Motion Discrimination Task

Four participants performed a direction discrimination task. On each trial, they viewed a patch of moving dots and decided whether the net motion was leftward or rightward. Task difficulty was manipulated by varying the probability that each dot was displaced in the coherent motion direction (left or right) when replotted 40 ms later, rather than being randomly repositioned (Roitman and Shadlen, 2002). Following standard terminology, we refer to this probability as *motion strength*.

Participants reported their response by moving a handle toward one of four targets corresponding to the joint reports: Left–High Confidence, Left–Low Confidence, Right–Low Confidence, and Right–High Confidence (Fig. 1A). The motion stimulus remained visible until a motor response was initiated, allowing choice, confidence, and response time (RT) to be recorded simultaneously from a single action.

**Figure 1.**
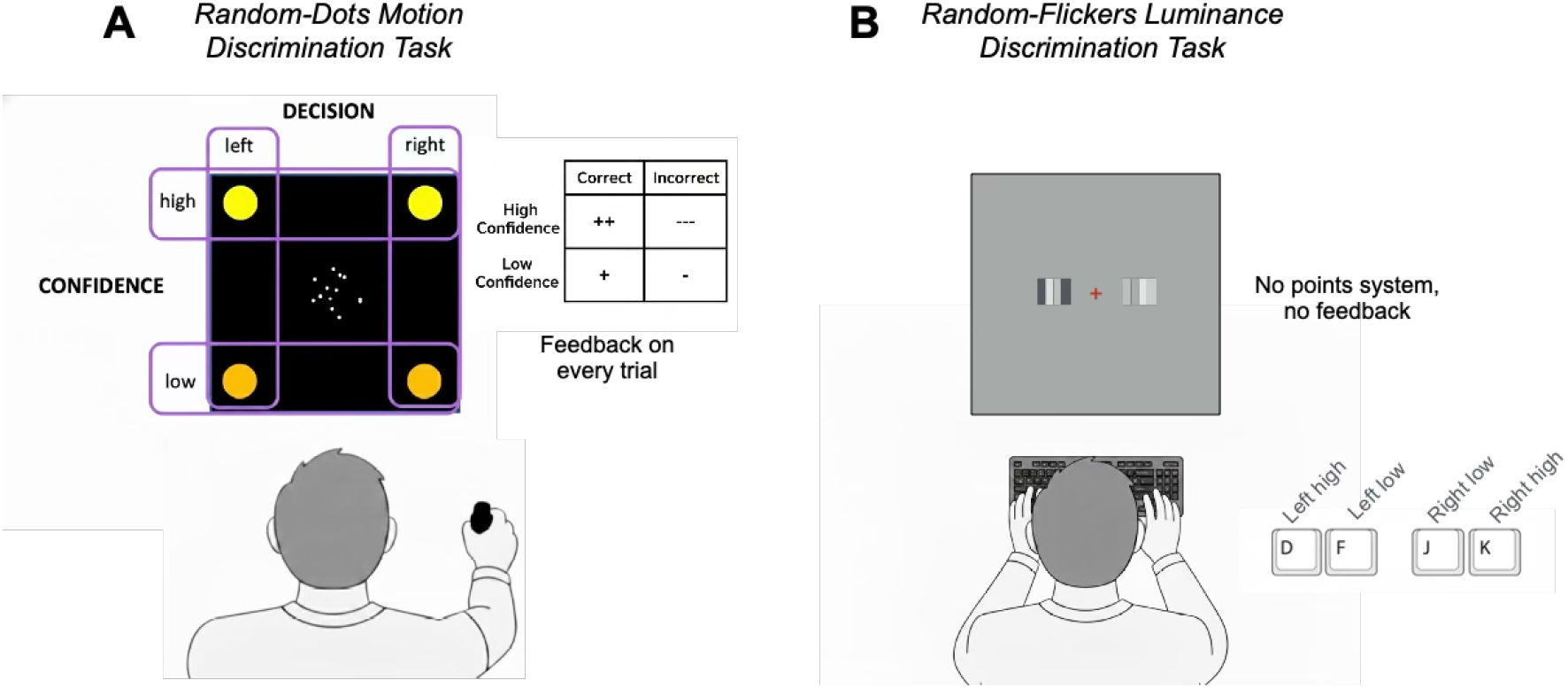
Motion and luminance discrimination with simultaneous choice-confidence reports. **(A)** Random-dots motion discrimination task. Participants performed a random-dot motion discrimination task, judging the direction of coherent motion (left or right). They reported their decision and confidence (low or high) by moving a handle constrained to a 2D plane toward one of four spatial targets (Left–High, Left–Low, Right–High, Right–Low). Confidence reports were incentivized through an asymmetric reward system where correct high-confidence choices earned +1 point but incorrect ones lost -1, while low-confidence choices earned +2 and lost -3 points. Feedback on accuracy and points earned was provided after every trial. The data from the motion task was published by van Den Berg et al. (2016). **(B)** Luminance discrimination task. Participants judged which of two patches was brighter (or darker, depending on the condition). Each patch consisted of four vertical bars. Choice and confidence were reported simultaneously using a computer keyboard: keys *d* and *f* indicated a leftward choice with high and low confidence, respectively, while keys *j* and *k* indicated a rightward choice with low and high confidence, respectively. . Unlike the motion task, confidence reports were not incentivized and feedback was not provided.

Confidence resolution was incentivized through an asymmetric payoff structure. Correct and incorrect low-confidence responses yielded +1 and −1 points, respectively, whereas correct and incorrect high-confidence responses yielded +2 and −3 points. After each decision, participants were informed of choice accuracy and the number of points earned. This dataset was analyzed in previous studies (Van den Berg et al., 2016; Zylberberg and Shadlen, 2025).

#### Luminance Discrimination Task

Four additional participants performed two versions of a luminance discrimination task. Participants fixated a central spot while two peripheral patches were presented simultaneously, one on each side of fixation, each consisting of four vertical bars (Fig. 1B) (Zylberberg et al., 2012; Mazor et al., 2023). In one version of the task, participants had to select the brighter patch (*Choose-Brighter*). In the other version, they had to select the darker patch (*Choose-Darker*). Because the decision is binary, the two formulations are logically equivalent.

Decision difficulty was manipulated by varying the nominal luminance difference between the two patches (termed *luminance strength*), and by corrupting the luminance of individual bars with additive zero-mean Gaussian noise that was updated every 66.7 ms. Participants reported their decision and confidence (high/low) simultaneously via keypress. As in the motion task, responses were self-paced and the stimulus terminated upon response. Unlike in the motion task, confidence reports were not incentivized and participants did not receive feedback about accuracy.

### Behavioral analysis

The relationship between choice, response time, and confidence is qualitatively similar across the motion and luminance tasks. The proportion of correct responses increased with stimulus strength (motion or luminance strength) (Fig. 2A)(Motion task: *β*_1_ = 21.92; Lum-choose-brighter: *β*_1_ = 71.55; Lum-choose-darker: *β*_1_ = 67.43; all *p* < 0.001; Eq. 21). This increase was more pronounced for decisions reported with high confidence (Mot: *β*_3_ = 2.46, *p* = 0.008; Lum-B: *β*_3_ = 29.57, *p* < 0.001; Lum-D: *β*_3_ = 13.14, *p* = 0.051). Decisions were faster for stronger stimuli, for both high- and low-confidence reports (Fig. 2B) (Mot: *β*_1_ = −1.65; Lum-B: *β*_1_ = −5.97; Lum-D: *β*_1_ = −5.56; all *p* < 0.001; Eq. 22). High-confidence decisions were overall faster than low-confidence decisions (Mot: *β*_2_ = −0.36; Lum-B: *β*_2_ = −0.57; Lum-D: *β*_2_ = −0.46; all *p* < 0.001), although this RT difference decreased as stimulus strength increased (Fig. 2B)(Mot: *β*_3_ = 0.56; Lum-B: *β*_3_ = 5.06; Lum-D: *β*_3_ = 4.27; all *p* < 0.001).

**Figure 2.**
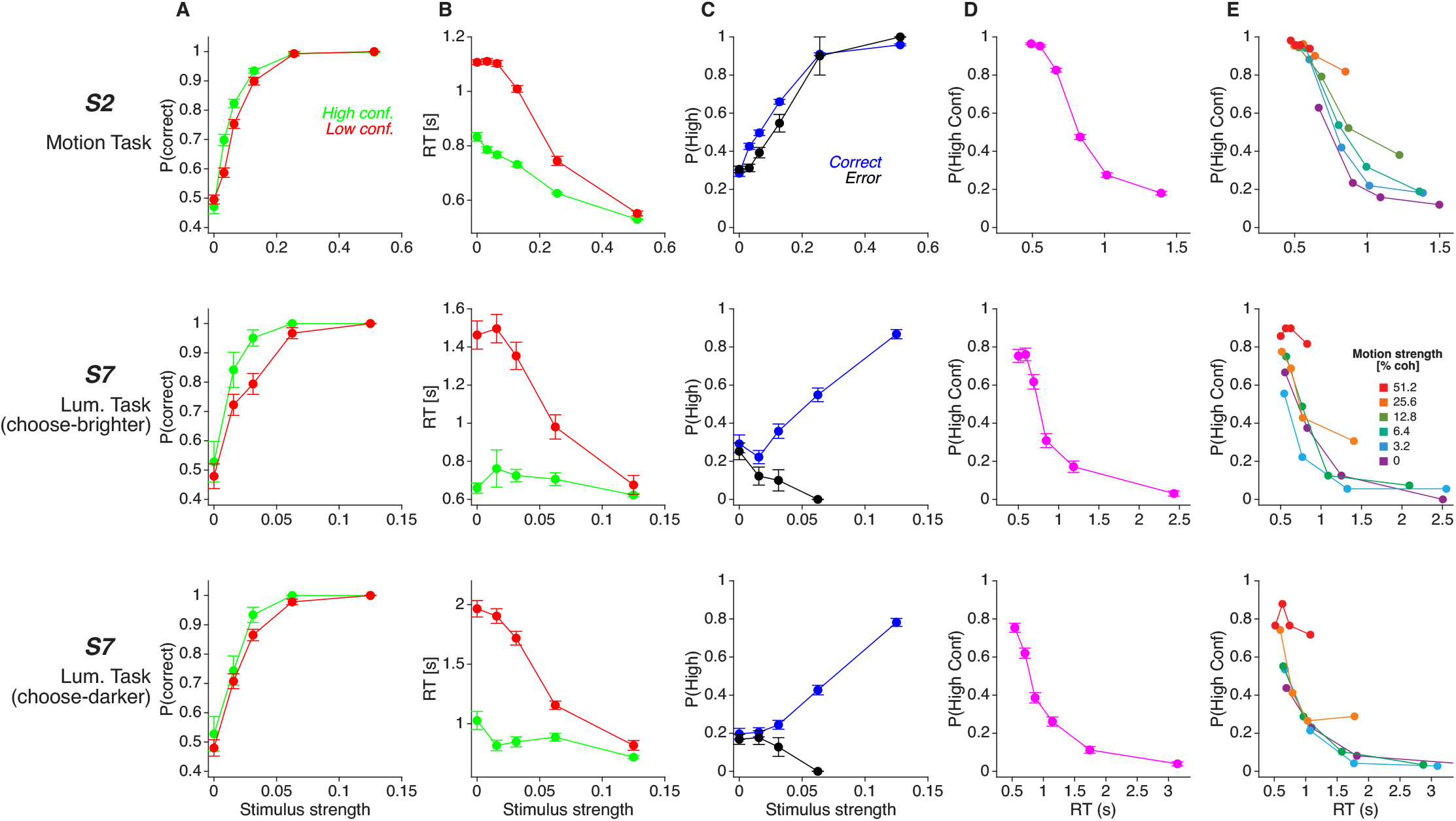
Behavior in the motion and luminance discrimination tasks. Behavioral data from representative participants performing the random dots motion task (top row) and the luminance discrimination task (Choose-Brighter variant: middle row; Choose-Darker variant: bottom row). **(A)** Proportion of correct choices for low-confidence (red) and high-confidence (green) reports as a function of stimulus strength (motion or luminance strength, depending on the task). For 0% stimulus strength the correct choice was assigned randomly. **(B)** Mean response time for low- and high-confidence reports as a function of stimulus strength. **(C)** Proportion of high-confidence responses as a function of stimulus strength, shown separately for correct and incorrect choices. **(D)** Proportion of high-confidence responses as a function of response time across all trials (correct and incorrect). Data were binned into six quantiles. **(E)** Proportion of high-confidence responses as a function of response time, separated by stimulus strength (correct choices only). Data were binned into four quantiles, independently for each level of stimulus strength. Data shown correspond to one representative participant per task; equivalent analyses for all participants are shown in subsequent figures. Error bars indicate s.e.m.

In the motion task, confidence increased with stimulus strength for both correct and incorrect responses (main effect of stimulus strength: *β*_1_ = 8.35, *p* < 0.001; interaction: *β*_3_ = 0.51, *p* = 0.548; Eq. 23), as has been shown previously for this dataset (van Den Berg et al., 2016). In contrast, in the luminance tasks, confidence increased with stimulus strength for correct decisions, but did not for incorrect decisions (Fig. 2C)(Lum-B: *β*_3_ = 35.22, *p* < 0.001; Lum-D: *β*_3_ = 21.11, *p* = 0.001; main effect of stimulus strength for incorrect decisions: Lum-B: *β*_1_ = −9.77, *p* = 0.250; Lum-D: *β*_1_ = 2.14, *p* = 0.730; Eq. 23).

In both tasks, confidence decreased markedly as RT increased (Fig. 2D) (Mot: *β*_1_ = −3.85; Lum-B: *β*_1_ = −3.04; Lum-D: *β*_1_ = −2.67; all *p* < 0.001). However, confidence was not solely determined by RT: even when restricting the analysis to correct trials, for a given RT confidence was higher at higher levels of stimulus strength (Fig. 2E) (Mot: *β*_2_ = 3.27; Lum-B: *β*_2_ = 19.66; Lum-D: *β*_2_ = 18.77; all *p* < 0.001; Eq. 24).

The average proportion of high-confidence responses was significantly higher in the motion task (0.76 ± 0.051 SEM) than in the luminance task (0.28 ± 0.045; *t*(6) = 7.00, *p* < .001) (Fig. 2C). This difference in confidence was observed despite no significant difference in decision accuracy, which was closely matched between the motion task (0.82±0.011) and the luminance task (0.81±0.012; *t*(6) = 0.55, *p* = .603). As we explain below, this difference in the proportion of high-confidence responses is critical for interpreting the model fits.

In summary, the relationship between choice, RT, and confidence is broadly similar across the two tasks, with two exceptions: (*i*) confidence increases with stimulus strength on error trials in the motion task, but not in the luminance task; and (*ii*) confidence is overall lower in the luminance task than in the motion task.

### Computational models

To adjudicate between the hierarchical and intentional accounts of confidence, we compared two computational models embodying these alternative architectures.

#### Hierarchical Model

In the hierarchical model, the choice is resolved by a competition between two drift–diffusion processes that integrate noisy sensory evidence over time (Fig. 3A-B). When one of the accumulators reaches the decision bound, the choice process terminates and a metacognitive readout mechanism is engaged. Confidence is derived from a two-dimensional confidence map that specifies how the probability of being correct depends on two variables: the state of the losing accumulator at decision time (the “losing race”) and the time taken to decide (Fig. 3C). The mapping from these variables to the probability that the chosen option is correct is obtained through Bayes’ rule, assuming a known prior distribution over motion strength and direction (Kiani et al., 2014). When the evidence difference between accumulators is large and the decision is reached quickly, the model assigns high confidence. Conversely, when the losing accumulator is close in value to the winning one, or when deliberation is prolonged, confidence decreases. Crucially, this architecture assumes that choice and confidence are functionally separable: the system first commits to an action and only then evaluates its certainty based on the state of the decision variables and the elapsed time.

**Figure 3.**
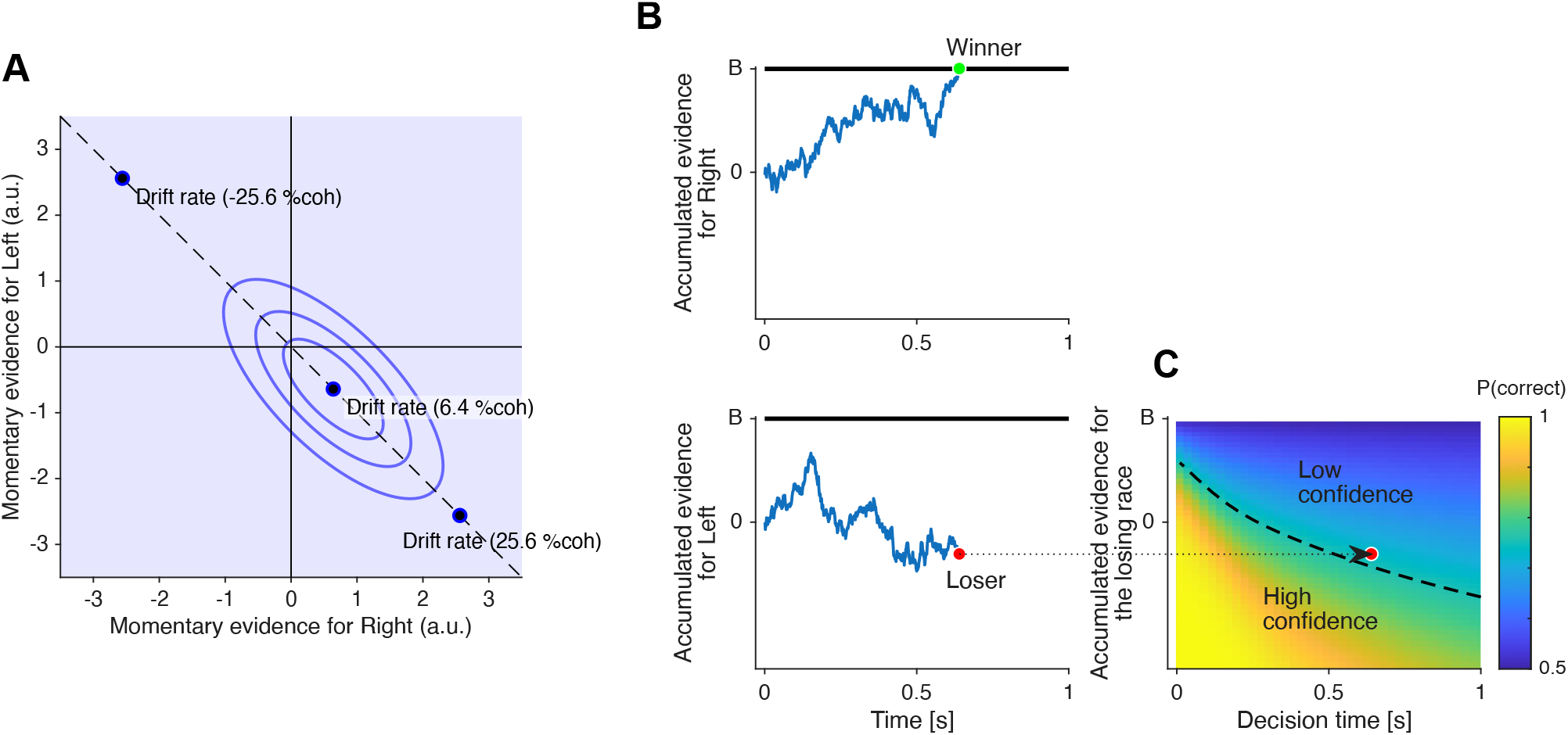
Hierarchical decision–confidence model. Schematic of a hierarchical architecture in which two competing drift–diffusion processes determine choice, and a separate readout mechanism computes confidence. **(A)** Distribution of momentary evidence for the two accumulators. Each sample provides evidence for the rightward and leftward alternatives. Blue contours denote iso-density levels of the joint distribution, highlighting the anti-correlated noise variability. Black dots indicate the mean drift rates for three example stimulus strengths. The degree of anti-correlation was treated as a free parameter and fit to the data. **(B)** Example trajectories of the two drift–diffusion processes over time. The decision is made when one accumulator reaches a fixed bound (*B*). **(C)** Two-dimensional confidence readout map. Confidence is computed by mapping the state of the losing accumulator (*x*_loser_) and the elapsed decision time (*t*) onto the posterior probability of being correct. Colors indicate this probability. A high-confidence report is generated when the probability exceeds a fixed threshold *θ*, partitioning the map into low- and high-confidence regions (dashed line).

#### Integrated (Intentional) Model

As an alternative to the hierarchical model, we implemented a ‘flat’ competitive architecture that collapses choice and confidence into a single intentional commitment. Rather than a two-stage process in which a decision is followed by a metacognitive readout, this account posits a simultaneous race between four integrated alternatives: “Left–High,” “Left–Low,” “Right–Low,” and “Right–High.” Our model is based on the RTCON model (Ratcliff and Starns, 2009, 2013).

The drift rate for each accumulator is determined by partitioning a latent evidence distribution (Fig. 4). At each moment, the stimulus generates a noisy sensory sample drawn from a Gaussian distribution. Three internal criteria (Fig. 4) divide this distribution into four regions. Each alternative is represented by its own accumulator, whose drift rate is proportional to the probability mass assigned to its corresponding region (Fig. 4). The decision process can be understood as if at each time step, a sample of sensory evidence is drawn and categorized according to the placement of the internal criteria (Fig. 4). The accumulator associated with the selected region receives a fixed increment (a “vote”) plus independent Gaussian noise, while the remaining accumulators accumulate only noise. For example, a strong rightward signal places most of its mass in the “Right–High Confidence” region, increasing the likelihood that this accumulator wins the race. A decision occurs when one of the four accumulators reaches its bound. Thus, choice, confidence and response time emerge jointly from a single competitive evidence-accumulation process.

**Figure 4.**
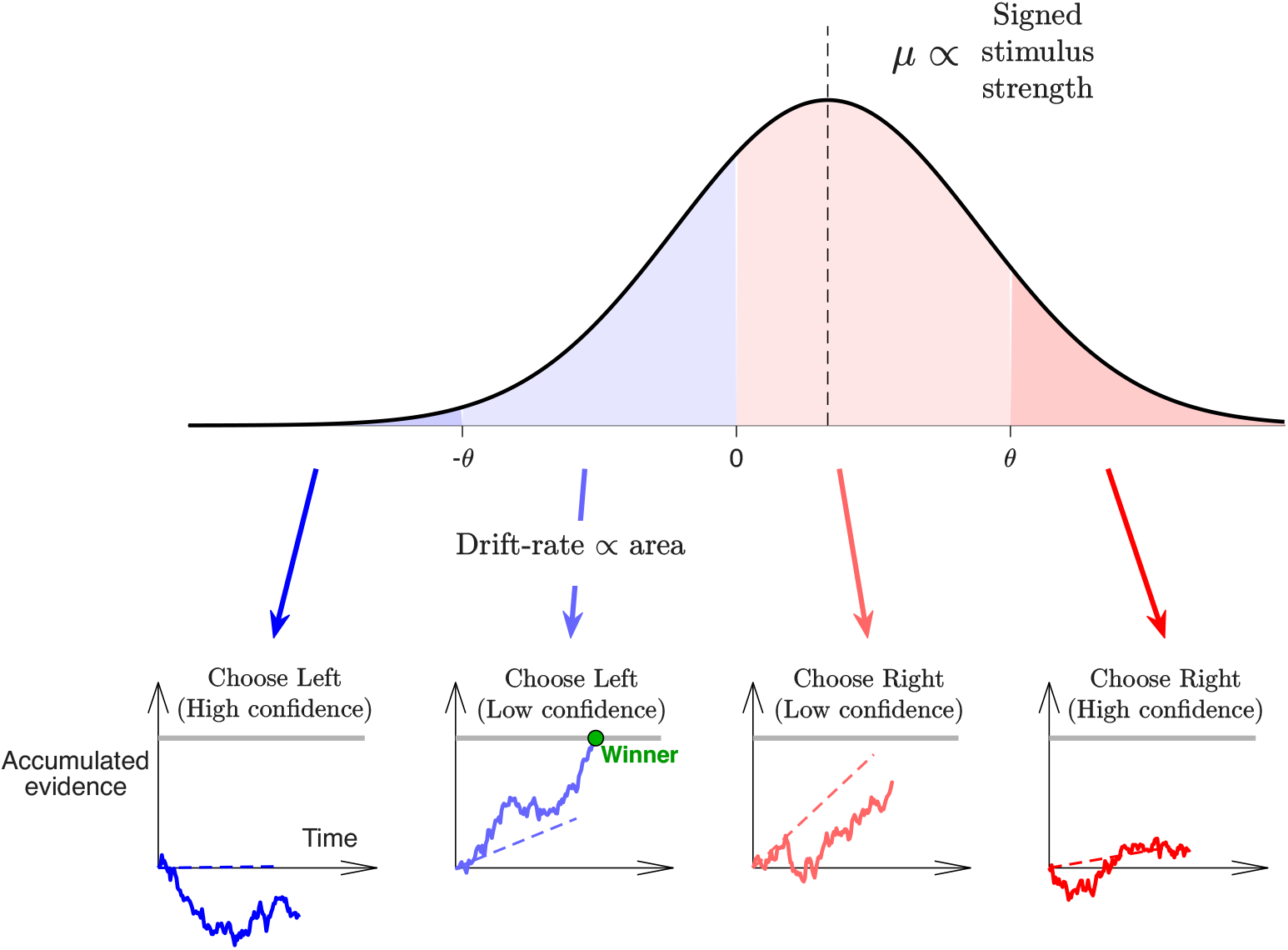
‘Flat’ competition model. **(top)** Evidence partitioning mechanism. In each time step, a sample of momentary evidence is drawn from a Gaussian distribution whose mean is proportional to the trial’s signed stimulus strength. This distribution is partitioned into four regions by three internal criteria (−*θ*, 0, *θ*). **(bottom)** The model consists of four drift-diffusion processes (races) that run in parallel: “Left-High”, “Left-Low”, “Right-Low”, and “Right-High”. The drift rate for each race is proportional to the area under the evidence distribution within its respective region. The races are independent conditioned on the trial’s signed stimulus strength. A decision is made as soon as one of the four accumulators reaches a upper bound, resolving choice and confidence simultaneously.

### Model comparison

To determine which architecture best accounts for the decision-making process, we fit both models to the data from the motion and luminance discrimination tasks. The models were fit to maximize the joint likelihood of choice, response time, and confidence on each trial (Methods).

#### Random dots motion Task

Both models successfully capture the central behavioral tendencies in the motion task (see Fig. 2), albeit with some limitations. The hierarchical model tends to overestimate the impact of confidence on accuracy, predicting poorer performance for low-confidence responses than is empirically observed (Fig. 5A). Additionally, the hierarchical model fails to reproduce the steep increase in confidence with motion strength on error trials (Fig. 5C). This mismatch occurs because the degree of anti-correlation between the evidence streams (*ρ*) in the best fitting model was relatively low: *ρ* = 0 for three of the participants and *ρ* = −0.5 for the remaining one. These low values of *ρ*, however, helped explain other aspects of the data. Indeed, constraining *ρ* = −0.7071—a value commonly assumed in prior work (Vivar-Lazo and Fetsch, 2025; van Den Berg et al., 2016; Kiani et al., 2014)—substantially degrades the model’s ability to account for response times in low-confidence decisions, revealing a trade-off in the hierarchical model’s ability to jointly account for the different aspects of the data (Fig. S1).

**Figure 5.**
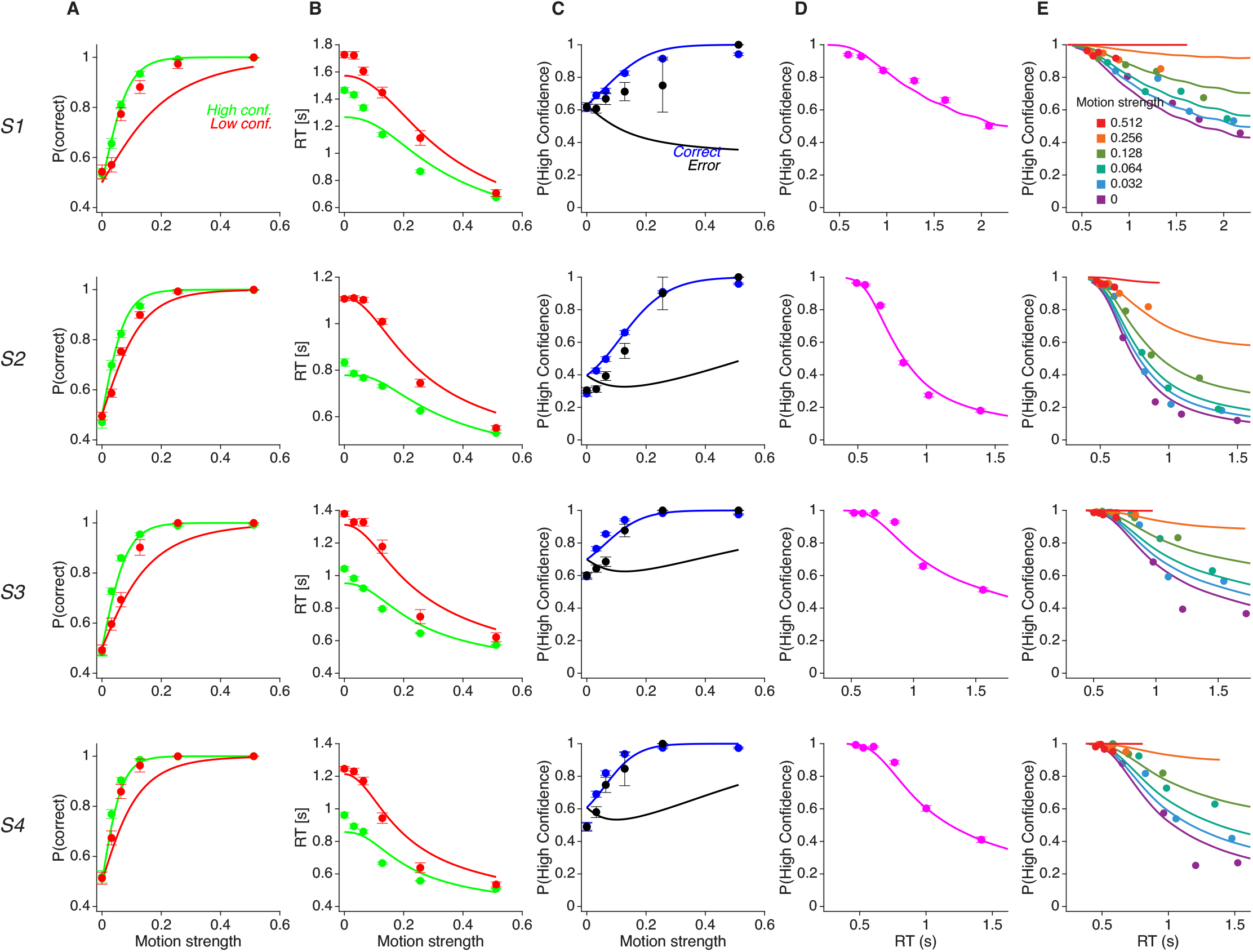
Behavior in the motion task and fits of the hierarchical model. **(A)** Proportion of correct motion choices for low-confidence (red) and high-confidence (green) reports as a function of motion strength. For 0% motion strength, the correct choice was assigned at random. **(B)** Mean response time for low- and high-confidence reports as a function of motion strength. **(C)** Proportion of high-confidence responses as a function of motion strength, shown separately for correct and incorrect choices. **(D)** Proportion of high-confidence responses as a function of response time. Trials were grouped into six quantiles. Both correct and incorrect choices were included. **(E)** Proportion of high-confidence responses as a function of response time, split by motion strength. Only correct choices were included. Each row corresponds to a different participant. Solid lines denote model fits. The dataset includes 5,245; 9,023; 9,021; and 4,999 trials for Participants 1–4, respectively. Error bars indicate s.e.m.

The flat model was also able to capture the association between choice, RT and confidence (Fig. 6). The model provides a superior account of the relationship between confidence and motion strength, capturing the strong positive correlation present in both correct and error trials (Fig. 6C). Surprisingly, the flat architecture also replicates Kiani et al. (2014)’s observation that, for a given RT, confidence is higher on trials with stronger motion (Fig. 6E). Because this relationship rules out models where confidence is strictly a time-dependent function, it has previously been used as evidence in favor of a hierarchical mechanism that weights decision time and the *balance-of-evidence*. However, our fits demonstrate that a single-stage competition is entirely sufficient to generate these dependencies without requiring a secondary evaluative process.

**Figure 6.**
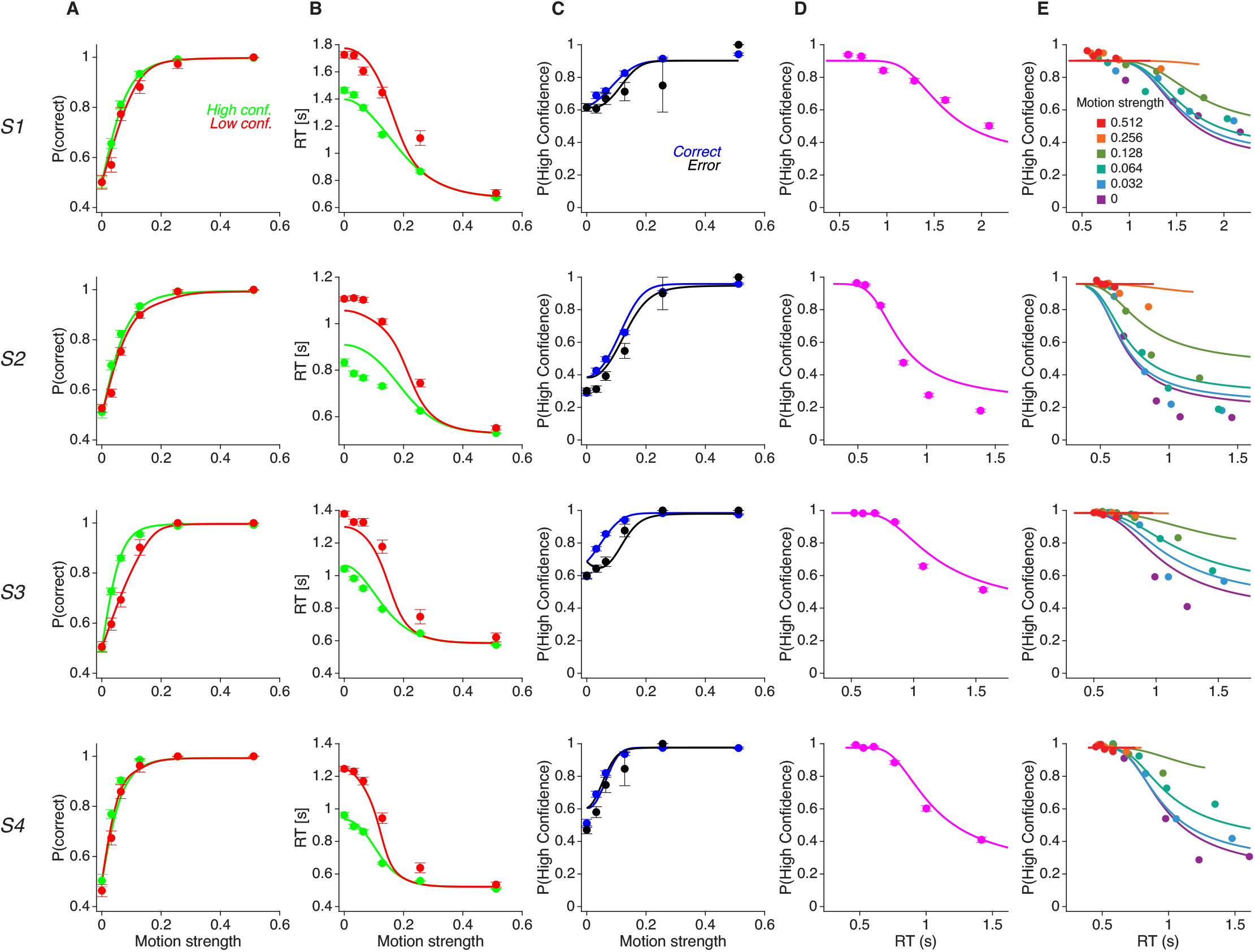
Behavior in the motion task and fits of the flat competition model. Same as Fig. 5 except that the solid lines represent fits of the ‘flat’ model.

Nevertheless, the integrated model fails to capture other aspects of the data. Most notably, for Subject 2 the model underestimates the difference in response times between high and low confidence decisions (Fig. 6B), and does not fully capture the decrease in confidence with response times (Fig. 6D).

To formally evaluate the two architectures, we computed the Bayesian Information Criterion (BIC) and Akaike Information Criterion (AIC). For three of the four participants, this quantitative comparison strongly favored the integrated model, while the hierarchical model provided a better fit for the remaining participant (Subject 2; Fig. 7, blue bars). In conclusion, while the flat model is imperfect, it outperforms the hierarchical model in the motion task. As we show in the following section, however, this advantage does not generalize to the luminance task.

**Figure 7.**
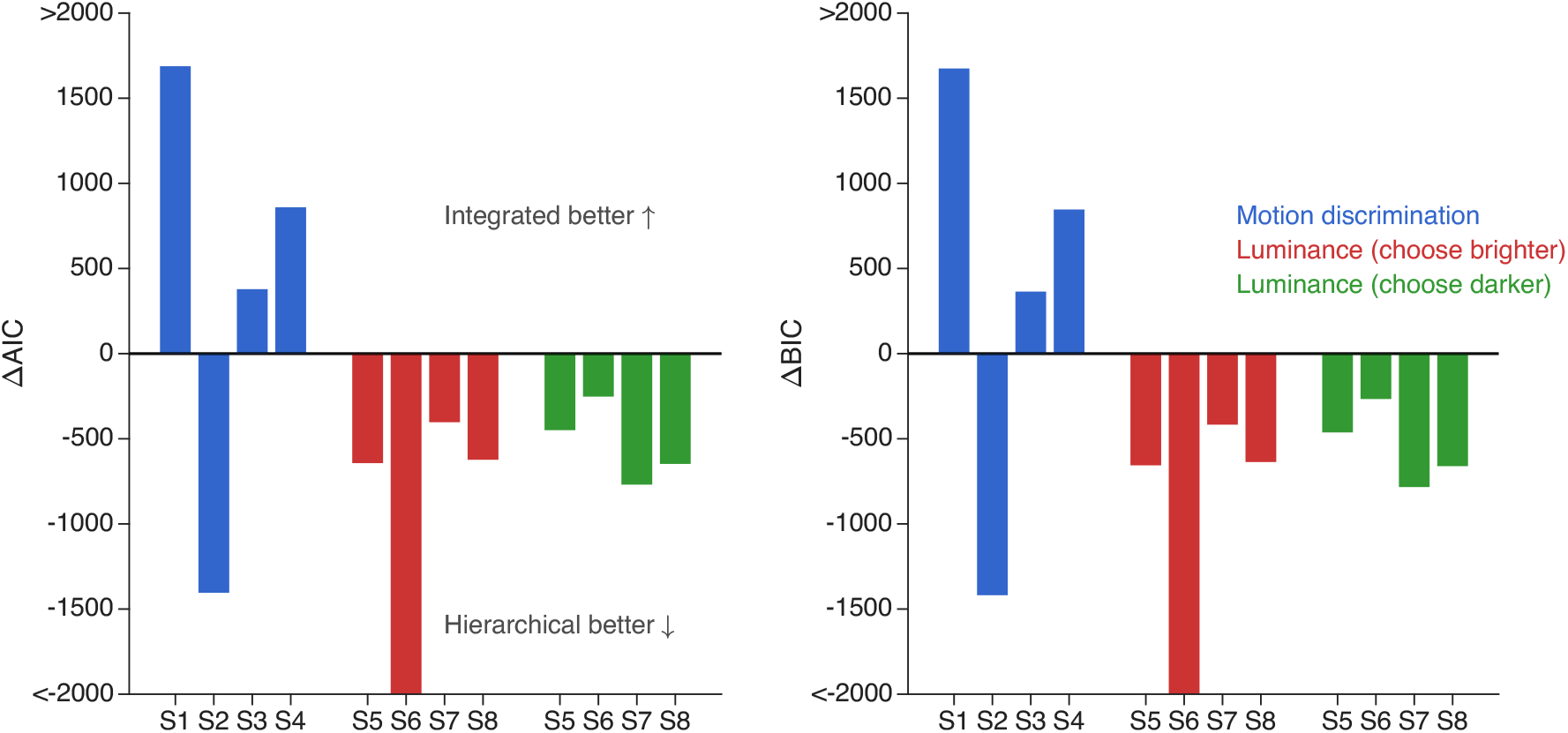
Model Comparison: AIC and BIC. Difference in Akaike (left) and Bayesian (right) Information Criteria between the hierarchical and flat competition models for the motion discrimination task (S1–S4) and the luminance tasks (S5–S8). Negative values indicate support for the hierarchical model, while positive values indicate support for the ‘flat’ model. In the motion task, the flat model provides a better fit for most participants. Conversely, in the luminance tasks (both “choose brighter” and “choose darker” variants), the hierarchical model is strongly favored.

#### Luminance-discrimination task

The hierarchical model achieved a good fit to the luminance task data, successfully capturing most empirical observations. Specifically, the model: (*i*) reproduced the observation that high-confidence decisions are faster and more accurate than low-confidence ones; (*ii*) captured the increase in confidence with luminance strength for correct choices, but not for incorrect choices; (*iii*) captured the overall association between confidence and response time (Fig. 8 and Fig. S2). In stark contrast, the flat model completely failed to fit the data (Fig. 9 and Fig. S3). Accordingly the model comparison using the AIC and BIC favored the hierarchical model for both versions of the task (Fig. 7).

**Figure 8.**
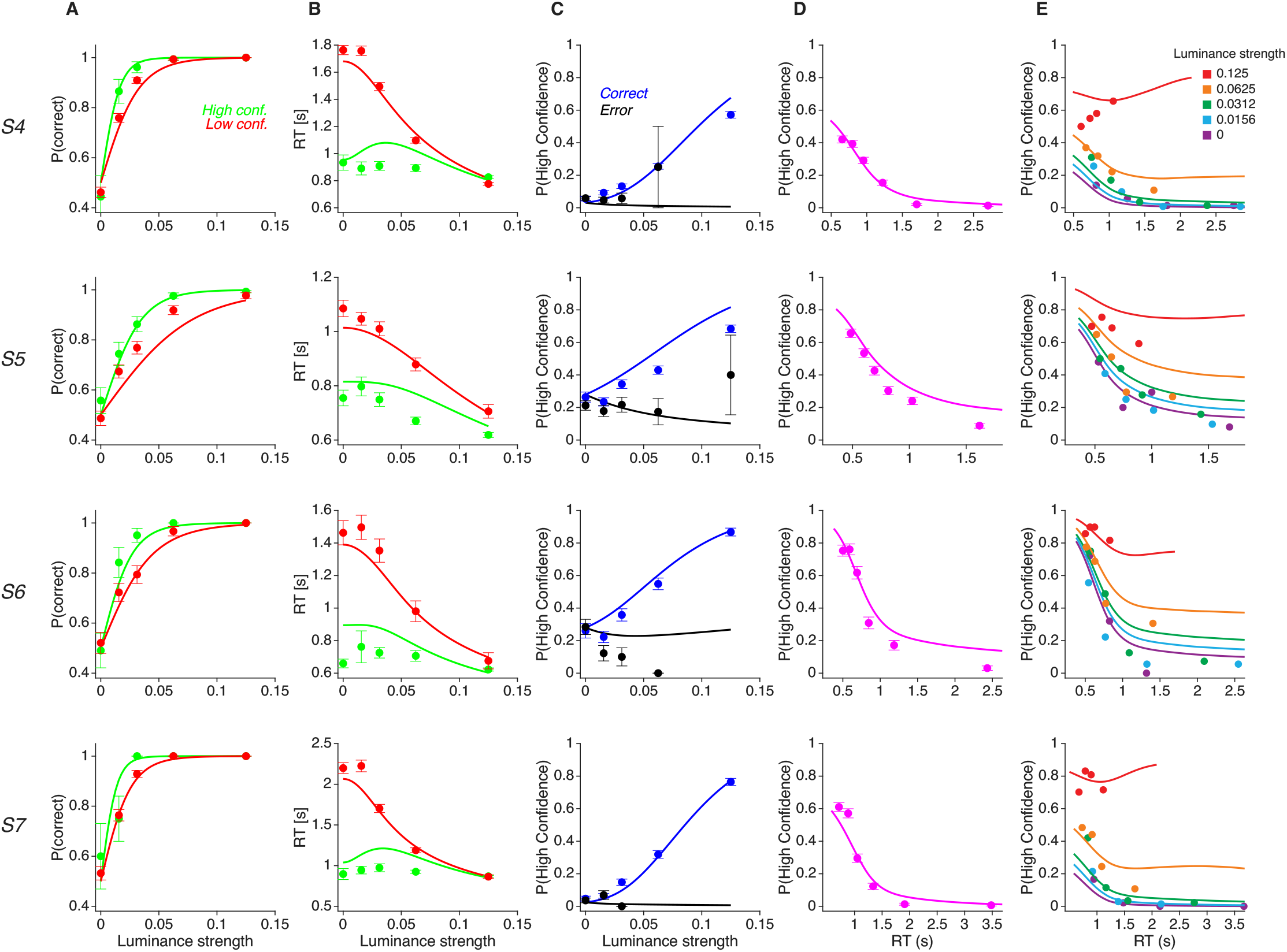
Behavior in the luminance (Choose-Brighter) task and fits of the hierarchical model. Same conventions as in Fig. 5.

**Figure 9.**
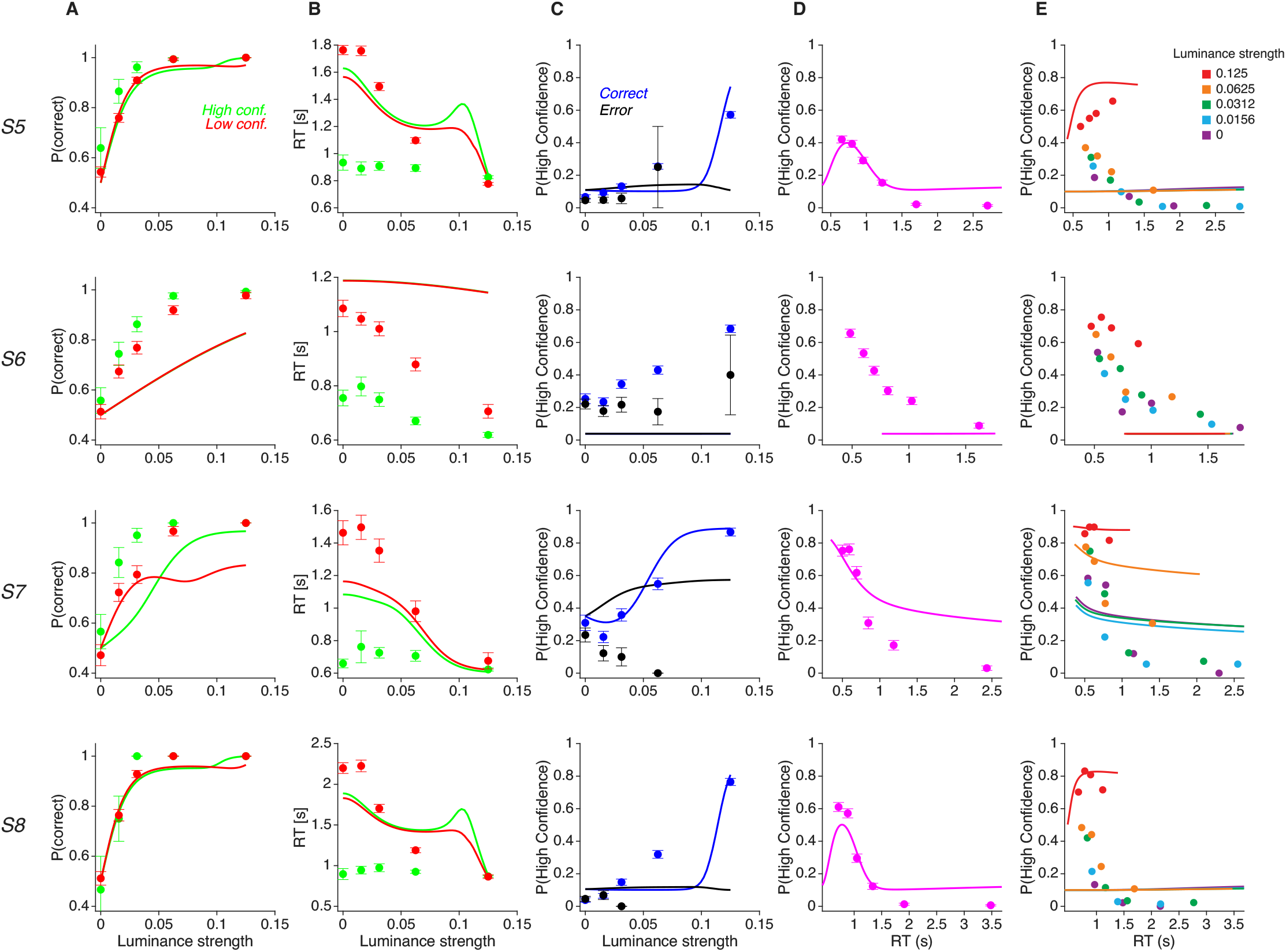
Behavior in the luminance (Choose-Brighter) task and fits of the flat competition model. Same conventions as in Fig. 5.

The failure of the flat model is surprising because, at a coarse level, the relationship between choice, RT, and confidence is similar in the luminance and the motion task, which the flat model successfully explains. The failure mode becomes clear when considering how the flat model generates high-confidence responses. Producing fast high-confidence choices requires that the corresponding accumulator either has a lower decision bound or a higher drift rate than its low-confidence counterpart. Both manipulations increase not only the speed of high-confidence responses, but also the probability that the high-confidence accumulator wins the race. As a result, a parameter regime that yields sufficiently fast high-confidence responses will predict a high overall frequency of such responses. While this pattern is consistent with the motion task, it directly contradicts the luminance data, where there are more low than high-confidence reports.

This tension reflects a structural limitation of the flat model: it intrinsically links the selection frequency of a confidence level to its associated response time and accuracy. Therefore, in its current form, the flat model cannot account for the pattern of behavior observed in the luminance task.

### Instruction-Dependent Positive-Evidence Bias

Although the hierarchical model substantially outperforms the flat model in the luminance task, it makes a simplifying assumption that the momentary evidence is computed as the instantaneous luminance difference between the two patches. This implies symmetric weighting, such that increasing the luminance of one patch by a certain amount has the same impact on behavior as decreasing the luminance of the other patch by the same amount.

Previous work has shown that this symmetry does not hold. Instead, fluctuations in the luminance of the selected patch exert a stronger influence on confidence than fluctuations in the unselected patch—a phenomenon often described as a positive evidence bias (PEB) (Zylberberg et al., 2012; Sepulveda et al., 2020; Peters et al., 2017; Rahnev and Denison, 2018). This asymmetry directly violates the model’s assumption of symmetric sensory weighting.

We confirmed this effect in the new dataset and extended it by showing that it depends on task instructions (Sepulveda et al., 2020; Mazor et al., 2023). In the Choose-Brighter task, the luminance of the selected patch was more informative about confidence than that of the non-selected patch (Fig. 10A; *β*_sel_ = 31.10, *β*_nsel_ = −17.75; Eq. 30); the test for symmetric weighting was highly significant (*H*_0_ : *β*_sel_ + *β*_nsel_ = 0, *p* < 10^−8^, Wald test).

**Figure 10.**
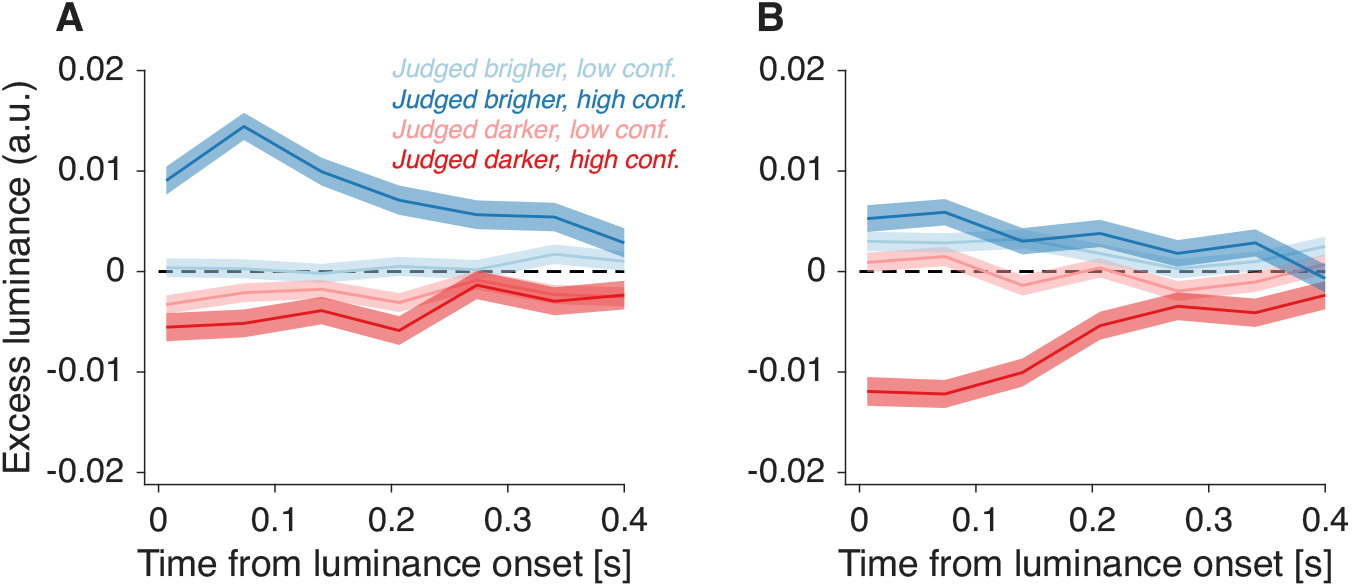
Psychophysical kernels of luminance fluctuations. **(A)** Choose-Brighter task. **(B)** Choose-Darker task. Traces show the time course of trial-averaged residual luminance fluctuations aligned to stimulus onset. Blue traces correspond to the patch judged brighter, and red traces to the patch judged darker. Line opacity indicates confidence level, with darker lines denoting high-confidence decisions and lighter lines denoting low confidence. Shaded regions represent ±1 standard error of the mean. Across both tasks, the separation between high- and low-confidence traces is most pronounced for the selected patch—i.e., the patch judged brighter in (A) and darker in (B)—consistent with an instruction-dependent positive evidence bias. Participants completed both task versions in counterbalanced order; analyses shown here include only trials prior to the task switch. Results including all trials are presented in Fig. S5.

Strikingly, the influence of luminance fluctuations on confidence depends on task instruction, despite both versions of the task being logically identical. In the Choose-Darker task, the same asymmetry was observed but with reversed sign, such that lower luminance in the selected patch was most informative about confidence (Figure 10B; *β*_sel_ = −32.31, *β*_nsel_ = 10.63; Eq. 30); the asymmetry test was again highly significant (*p* < 10^−8^, Wald test).

These asymmetric and instruction-dependent kernels demonstrate that the sensory decision variable cannot be reduced to a simple luminance-difference computation. However, this limitation concerns the construction of the low-level sensory variable rather than the overarching computational architecture linking choice, response time, and confidence. The central conclusion of the model comparison therefore remains: the joint relationship between choice, RT, and confidence requires a computational architecture in which metacognitive evaluation occurs in a distinct, hierarchical stage. The flat model fails at this architectural level, whereas the hierarchical model succeeds despite its simplified assumption about low-level sensory weighting.

## Discussion

We contrasted probabilistic and intentional approaches to computing confidence in decision-making. In the probabilistic framework, confidence is computed as the posterior probability of being correct given the evidence and time, whereas in the intentional framework, the actions associated with each possible choice-confidence combination compete until the accumulated evidence in favor of one crosses a threshold. We examined how representative models of these two approaches account for choice, response time, and confidence in two perceptual tasks: a motion discrimination task, in which confidence was incentivized through a reward structure favoring high-confidence decisions when they were more likely to be correct, and a luminance discrimination task, in which confidence was not incentivized. Neither model emerged as clearly superior: the intentional model provided a better account of behavior in the motion task, whereas the hierarchical model more accurately captured performance in the luminance task.

This divergence may reflect task-dependent shifts in cognitive strategy driven by external reward contingencies. The motion task, with its point system and trial-by-trial feedback, may have conditioned participants to treat the four choice-confidence combinations as a ‘flat’, integrated competition. In contrast, in the unincentivized luminance task, the brain may rely on a hierarchical architecture, generating confidence through a second-order metacognitive computation. Alternative explanations cannot be ruled out, such as differences in the stimuli (motion versus luminance) or variations in training and experience. Nevertheless, these findings underscore the flexibility and diversity of the mechanisms underlying confidence computation across contexts.

Hierarchical and integrated strategies imply markedly different neurophysiological implementations of confidence computation. The hierarchical model involves a sequence of complex operations, as the brain must access the state of the losing decision variable, register decision time, estimate the probability of being correct given these variables, and then compare this estimate to a threshold to determine whether to report high or low confidence. All of this must occur rapidly, as response time in this model is determined by the evidence accumulation process that precedes confidence computation (van Den Berg et al., 2016; Kiani et al., 2014; Vivar-Lazo and Fetsch, 2025). Although some components could, in principle, be computed in parallel with the decision itself, the overall procedure remains computationally demanding and relies on neural mechanisms that are not yet understood. In contrast, the integrated model relies on computations similar in complexity to those required for a standard multi-alternative decision process (Churchland et al., 2008). Evidence accumulates directly in action representations associated with specific choice–confidence combinations, suggesting that the competition manifests in brain areas involved in action preparation (Shadlen et al., 2008).

Recent neurophysiological evidence in non-human primates supports the integrated model’s prediction that the unfolding decision process is observable in action-preparation circuits. Vivar-Lazo and Fetsch (2025) trained macaques on a motion discrimination task similar to ours, incentivizing confidence with asymmetrical rewards. Monkeys reported both choice and confidence simultaneously via saccadic eye movements toward distinct targets. In the lateral intraparietal cortex (LIP), neurons encoding the intention to direct gaze toward each target (*T*_*in*_ neurons) exhibited joint selectivity for choice and confidence approximately 200 ms after motion onset, and their activity displayed characteristics of a drift-diffusion process, as predicted by the integrated model. By contrast, the hierarchical model provides no clear predictions for the activity of *T*_*in*_ neurons: while the hierarchical model posits that evidence accumulation occurs in a two-dimensional space (e.g., left versus right), the *T*_*in*_ neurons encode specific combinations of choice and confidence. Therefore, if confidence is computed only at decision commitment, as the hierarchical model proposes, it is unclear why *T*_*in*_ neurons would show drift-diffusion-like dynamics during deliberation (Vivar-Lazo and Fetsch, 2025; Vivar Lazo, 2024). Since the integrated model better captures behavior in our task, we hypothesize that monkeys solve it through an integrated strategy, although this requires formal evaluation using the data from Vivar-Lazo and Fetsch (2025).

Our study is subject to several important limitations. First, while we categorized models into hierarchical and integrated classes, we only evaluated a single representative implementation from each. In particular, the space of integrated models is broader than the specific implementation considered here, and may include variants with flexible confidence criteria, stimulus-dependent variability, or interactions between accumulators, which could give rise to richer behavioral patterns (Ratcliff and Starns, 2009, 2013). It is therefore conceivable that a variant of the integrated model might explain the luminance task data more effectively than the current hierarchical model, and thus we should avoid overgeneralizing these results to all architectures within each class. This is a common hurdle in model comparison (Zylberberg and Shadlen, 2016; Latimer et al., 2015).

Participants in the luminance task exhibit an instruction-dependent positive evidence bias (PEB) (Zylberberg et al., 2012), which demonstrates that Choose-Brighter and Choose-Darker instructions, though logically identical, lead to different decision strategies (Sepulveda et al., 2020). Our hierarchical model, based on the Balance-of-Evidence (BoE) hypothesis, does not capture this effect. Potential modifications to the model to account for this bias include attentional asymmetries, or variations in the weighting or processing speed of the evidence in favor of each choice alternative (Sepulveda et al., 2020; Zylberberg et al., 2012; Paz et al., 2016). Additional evidence against the BoE is derived from the analysis of neural data. While the BoE posits that confidence depends on the state of the losing accumulator, recent work shows that the accumulator for the chosen option contains significantly more confidence-related information than the unchosen one (Zylberberg and Shadlen, 2025; Toso et al., 2025). A comprehensive model integrating these neural and behavioral complexities has yet to be developed.

In turn, the numerous reported dissociations between choice and confidence pose a challenge to the generality of integrated models of confidence (see, for example, Fleming and Daw (2017); Rahnev (2026) for a review). These dissociations may arise from manipulations of the stimulus and attention (Wilimzig et al., 2008; Vlassova et al., 2014; Rahnev et al., 2011; Lau and Passingham, 2006; Graziano and Sigman, 2009; Bona and Silvanto, 2014; Zylberberg et al., 2012), differences in brain structure (Fleming et al., 2010; Baird et al., 2015; Barttfeld et al., 2013), brain lesions (Del Cul et al., 2009; Fleming et al., 2014; Lak et al., 2014; David et al., 2012; Komura et al., 2013; Rounis et al., 2010), and individual differences (Ais et al., 2016; Song et al., 2011; Palmer et al., 2014). It is relatively straightforward to conceive how such dissociations could arise in a hierarchical model—for example, through interference with the readout process or by degrading the information on which the metacognitive judgment is based. In contrast, it is less clear how integrated models can reproduce these effects, since biases affecting confidence would be expected to also influence the primary decision.

Taken together, these observations suggest possible limits to the generality of both integrated and hierarchical models and raise the possibility that the computation of confidence may depend on task features such as confidence incentives, trial-by-trial feedback, and whether confidence is reported sequentially or simultaneously with the choice. These predictions could be tested empirically (e.g., by independently manipulating incentives).

The question we aim to address in this study can be framed within a broader question: to what extent must problems that possess a hierarchical structure at the computational level be implemented hierarchically at the algorithmic level in the brain? This issue has been debated in the domain of sequential perceptual choices; for instance, behavior initially thought to require independent, serial sub-decisions (Lorteije et al., 2015) may be equally well explained by a ‘flat’ competition among the final motor actions, driven by mutual inhibition (Hyafil and Moreno-Bote, 2017; Zylberberg et al., 2017). By contrasting a hierarchical model with an integrated one for simultaneous choice-confidence reports, we highlight the possibility that the brain may, under the right conditions, transform a second-order metacognitive evaluation into a flat, first-order competition among actions.

## Methods

### Random dots motion discrimination task

Four participants completed a motion direction discrimination task in which they judged whether the net motion of a random-dot stimulus was leftward or rightward. Trials with leftward and rightward motion were randomly interleaved. Motion strength—defined as the probability that a dot was displaced coherently in the motion direction rather than randomly replotted—varied pseudo-randomly across trials and took one of six values: 0%, 3.2%, 6.4%, 12.8%, 25.6%, or 51.2%. We refer to signed motion strength (negative for leftward motion) as *motion coherence*. On 0% coherence trials, the correct response was assigned randomly. Stimulus-viewing duration was controlled by the participants. When ready, participants reported both their choice and confidence (high vs. low) simultaneously by moving the handle of a robotic manipulandum (Howard et al., 2009) toward one of four targets. The targets were arranged such that horizontal position encoded choice (left vs. right) and vertical position encoded confidence (high vs. low). High- and low-confidence targets were positioned at the top and bottom of the screen, respectively, with their mapping reversed across blocks to counterbalance potential spatial biases. Additional experimental details are provided in the original publication (van Den Berg et al., 2016).

### Luminance discrimination task

Four additional participants (20–30 years old) completed a reaction-time luminance discrimination task in which they compared the luminance of two patches presented simultaneously to the left and right of a central fixation spot. Depending on the session, participants were instructed to report either the brighter or the darker patch. Because the decision is binary, the two formulations are logically equivalent.

Choice and confidence were reported simultaneously via a computer keyboard: the “d” and “f” keys indicated a leftward choice with high and low confidence, respectively, whereas the “j” and “k” keys indicated a rightward choice with low and high confidence, respectively. Stimuli remained visible until the participant made a response. Unlike in the motion task, confidence reports were not incentivized and participants did not receive feedback about choice accuracy.

#### Luminance stimuli

The luminance stimuli consisted of two patches presented symmetrically at an eccentricity of 1^◦^ relative to the center of each patch. Each patch was composed of a 4 × 1 grid of small rectangles (vertical bars), with each individual rectangle measuring 0.56^◦^ in height and 0.14^◦^ in width. The patches were displayed against a uniform gray background with a baseline intensity (*I*_*bg*_) of 0.5 (i.e., 50% of the maximum luminance of the monitor). Given a maximum monitor luminance of 36 cd/m^2^ (gamma corrected), this background was maintained at 18 cd/m^2^.

To manipulate task difficulty, the mean luminance difference between the two patches (termed luminance strength, Δ*L*) was varied across trials using five levels: 0, 0.0156, 0.0312, 0.0625, and 0.1250. We define the *signed luminance strength* as Δ*L*, taking negative values when the left patch was brighter. On each trial *i* for participant *j*, the mean generative luminance for a given patch *p* ∈ {*L, R*} was defined as:

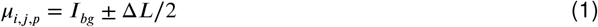

To enable psychophysical reverse correlation, the luminance of individual bars was corrupted with additive zero-mean Gaussian noise. Specifically, the luminance of each of the four bars within a patch was independently sampled from a normal distribution:

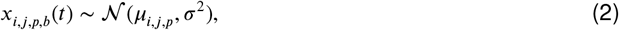

where *b* ∈ {1, …, 4} indexes the bars within a patch, *t* indexes the time samples, and *σ* = 0.1 is the standard deviation of the stimulus noise. These luminance values were updated synchronously every 5 frames—66.7 ms at the 75 Hz monitor refresh rate—to create a dynamic flickering effect.

#### Choose-Brighter and Choose-Darker variants

Participants S5-8 completed 3114, 2042, 974 and 1849 of the Choose-Brighter task and 3997, 1948, 1974 and 1967 trials of the Choose-Darker task, respectively, in counterbalanced order. In each session, participants performed only one of the two versions of the task, in order to minimize interference between the two sets of instructions. Fig. S4 shows, for the four participants, the order in which they completed the trials for each variant of the experiment.

#### Ethics Statement

The study was conducted at Columbia University (New York). All participants provided written informed consent. The study was approved by the Institutional Review Board of Columbia University Medical Center.

### Hierarchical model

The decision process is modeled as a 2D Correlated Drift-Diffusion Model (DDM) where two accumulators race toward an upper decision bound *B*. The state of the two accumulators, **x**(*t*) = [*x*_1_(*t*), *x*_2_(*t*)]^*T*^, evolves according to a stochastic differential equation:

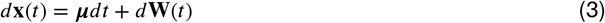

The drift vector ***μ*** is determined by the signed stimulus strength (*C*), a sensitivity parameter (*κ*), an urgency signal (*k*_urg_), and a subjective bias (*C*_0_). The urgency and bias parameters are explicitly scaled into units of stimulus strength by multiplying them by *κ*. This yields the specific drift rates for the two choices (Right and Left):

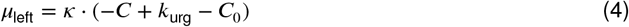

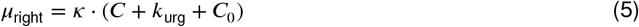

The noise term *d***W**(*t*) is a bivariate Wiener process with a covariance matrix Σ. The correlation coefficient *ρ* between the two evidence streams is constrained by the parameter *k*_*solv*_ (an integer ≥ 2) to allow for an analytical solution via the Method of Images:

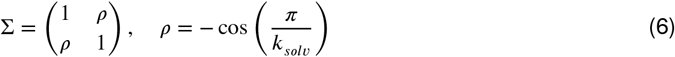

A decision is registered when either accumulator reaches *B*, and the predicted response time is calculated as the bound-crossing time *t* plus a non-decision time *T*_*nd*_.

Confidence is calculated as the posterior probability that the choice is correct, evaluated using the state of the losing accumulator (*x*_*loser*_) at the decision time *t*. Because the true signed stimulus strength is unknown to the subject, the model computes confidence maps by marginalizing over its possible values using a prior distribution *P* (*C*). For a given choice (e.g., “Right”), the confidence map is defined as:

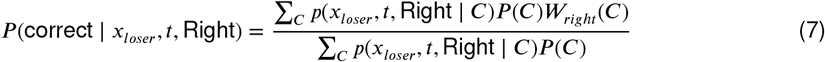

where *p*(*x*_*loser*_, *t* ∣ Right, *C*) is the spatial density of the losing accumulator extracted via the Method of Images flux (Shan et al., 2019). The prior *P* (*C*) is uniform over the discrete set of values of *C* used in the motion or luminance experiment (with double weight assigned to *C* = 0), and the choice-congruent weight *W*_*right*_(*C*) is set to 1 for *C* > 0, 0.5 for *C* = 0, and 0 for *C* < 0.

The model applies a confidence criterion parameter (*θ*) to segment the continuous probability map into a binary high/low confidence mask. For the ‘Right’ choice, the high-confidence mask *M*_*high*_ is:

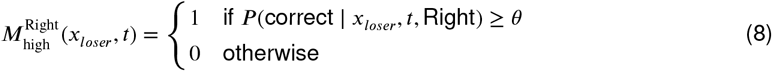

To compute the likelihood of observing a specific confidence report at a given decision time *t* and signed stimulus strength *C*, the model integrates the probability density of the losing accumulator over the spatial bins defined by this mask:

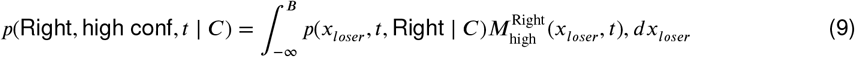

In practice, this is computed as the discrete sum over the spatial grid arrays at the interpolated empirical decision time (*t* = *RT* − *T*_*nd*_).

The model optimizes 7 free parameters: **Θ** = {*κ, B, T*_*nd*_, *θ, k*_urg_, *C*_0_, *k*_*solv*_}. The best-fitting parameter values are shown in Table S1. Parameters are estimated by minimizing the negative log-likelihood (NLL) of the joint distribution over all trials *i*:

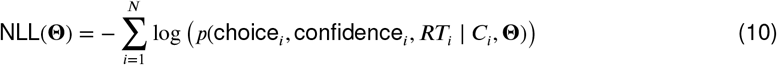

Our implementation of the model closely resembles prior ones (Kiani et al., 2014), including one used to analyze the same motion task dataset we analyzed here (van Den Berg et al., 2016). Our implementation differs in two key respects: (*i*) the degree of noise anticorrelation between the evidence streams feeding the two accumulators is treated as a free parameter rather than being fixed a priori; (*ii*) we fit the data by maximizing the joint likelihood of choice, response time, and confidence, in contrast with previous implementations of the model that either fit RT alone (Kiani et al., 2014), fit summary statistics (van Den Berg et al., 2016), or allowed confidence to become decoupled from the probability of being correct at long response times (Vivar-Lazo and Fetsch, 2025).

### Integrated Competition Model

The integrated (‘flat’) model was based on the RTCON model (Ratcliff and Starns, 2009). The model comprises four competing drift-diffusion processes, each one corresponding to a unique combination of choice (Left/Right) and confidence (High/Low).

We assume that the motion or luminance stimulus gives rise to an evidence distribution that is normally distributed with unitary variance. The effective mean of this distribution is determined by the motion coherence (or signed luminance strength), *C*, a sensitivity scaling parameter, *κ*, and a subjective bias, *C*_0_:

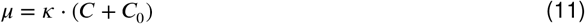

The drift-rate for each alternative is obtained partitioning the evidence distribution by a confidence criterion (*θ*). The probability mass areas of the normal distribution 𝒩 (*μ*, 1) bounded by [−*θ*, 0, *θ*] define the relative strengths of the four drift-diffusion processes:

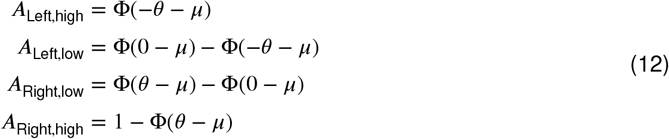

where Φ is the cumulative distribution function of the standard normal distribution. The final drift rate *v*_*k*_ for race *k* is scaled by a maximum rate *v*_*max,k*_:

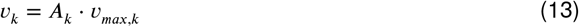

The maximum rate is *v*_*max,l*_ for low-confidence decisions and *v*_*max,h*_ for high-confidence ones.

The decision process terminates when one of the four accumulators reach an upper bound *B*_*k*_. The boundaries are *B*_*h*_ for high-confidence races and *B*_*l*_ for low-confidence races. The time to reach a bound for a single race follows a Wald (Inverse Gaussian) distribution. For a given race *k* with drift *v*_*k*_ and decision bound *B*_*k*_, the probability density function (*f*) and the survival function (*S*) at time *t* (where *t* = *RT* − *T*_*nd*_ after subtraction of the non-decision time) are:

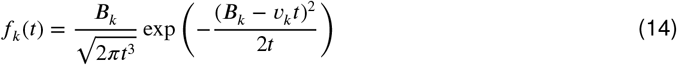

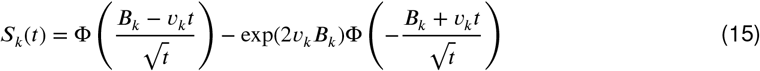

In a race between *N* = 4 accumulators, the probability that channel *k* wins at time *t* is the probability that it reaches its bound at *t* while all other channels have not yet reached theirs:

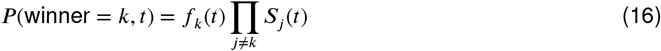

To account for lapses or errors that may occur after choice commitment, we include a “flip” parameter *p*_flip_, representing the probability that a subject reports the opposite confidence level than the one dictated by the winning accumulator. Thus, for a Leftward High-confidence response (Left_high_):

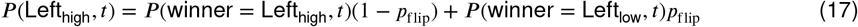

The total log-likelihood for a dataset is the sum of the log-probabilities of the observed response (choice, confidence) and RT for every trial *i*:

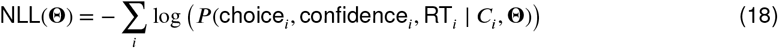

where the free parameters are **Θ** = {*θ, κ, C*_0_, *B*_*h*_, *B*_*l*_, *v*_*max,l*_, *v*_*max,h*_, *p*_flip_, *T*_*nd*_}. The best-fitting parameter values are shown in Table S2.

### Modeling the Choose-Darker task

To fit the data from the Choose-Darker version of the luminance task, we applied the following transformation. We recoded each response into the equivalent report that would have been obtained had participants been performing the Choose-Brighter version. For example, if a participant selected *right* with high confidence in the Choose-Darker task, we modeled this response as if the participant had selected *left* with high confidence in the Choose-Brighter task. This allows us to use the same model for the Choose-Brighter and Choose-Darker versions. Computationally, this transformation is equivalent to multiplying the signed luminance strength by −1 while keeping the original choice unchanged. Although it is clear from Fig. 10 that that instruction influences how the task is solved, the models considered in this study do not have the capacity to accommodate this distinction and instead treat the two versions of the task as equivalent.

### Model fitting

The models were fit to each participant’s single-trial data. Maximum likelihood estimation was performed using the BADS algorithm (Acerbi and Ma, 2017), minimizing the negative log-likelihood of the joint distribution of choice, confidence, and RT. We initialized the optimization from multiple (N = 10) randomly selected starting points within plausible parameter bounds to minimize the risk of convergence to local minima.

### Model Comparison

We assessed model parsimony and goodness-of-fit using AIC and BIC, which penalize the maximum likelihood for the number of free parameters to prevent overfitting. For each model *m*, these were calculated as:

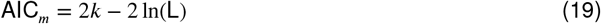

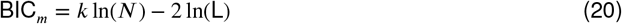

where *k* represents the number of free parameters, L is the maximum likelihood estimate, and *N* is the total number of trials per subject. Lower values of AIC and BIC indicate a superior model.

### Data analysis

Single-trial behavioral data were analyzed using mixed-effects regression models to account for the hierarchical structure of the data. In all models described below, *i* indexes the individual trial and *j* indexes the participant. We define evidence, *E*_*ij*_, as either the motion or luminance strength. Response times (RT_*ij*_) were log-transformed prior to analysis to mitigate the characteristic positive skew of RT distributions and were analyzed using mixed-effects linear regression. Binary dependent variables, including choice accuracy (*A*_*ij*_ ∈ {0, 1} for incorrect/correct) and confidence (conf_*ij*_ ∈ {0, 1} for low/high), were analyzed using mixed-effects logistic regression.

To test how stimulus strength and confidence interact to predict choice accuracy, we fit a logistic regression model defined as:

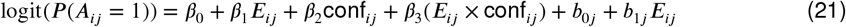

Similarly, to test how decision speed scales with stimulus strength for both high- and low-confidence reports, log-transformed RTs were modeled using a linear regression defined as:

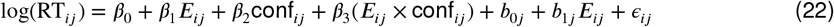

We also used logistic regression to evaluate the predictors of high versus low confidence. To test for a divergence in how stimulus strength influences confidence on correct versus error trials, we fit:

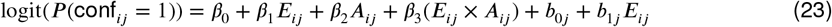

Finally, to determine if stimulus strength provided explanatory variance for confidence independent of decision time, we restricted the analysis to correct trials and fit a logistic regression predicting confidence from both RT and evidence:

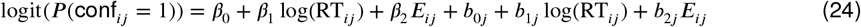

In all equations, the *β* terms represent the fixed effects estimating the overall group-level relationships, and the *b* terms represent the subject-level random effects (participant-specific variations), assumed to be normally distributed with a mean of zero. For continuous models, *ϵ*_*ij*_ represents the trial-level residual error. For each fixed effect, we report the group-level coefficient estimate (*β*) and the associated *p*-value, which were derived using *t*-tests for the linear models and Wald *z*-tests for the logistic models.

### Confidence kernels

To estimate the influence of stochastic stimulus fluctuations on confidence, we computed psychophysical kernels using the frame-by-frame luminance values defined previously (Eqs. 1 and 2). We first average the luminance of the four bars in each patch:

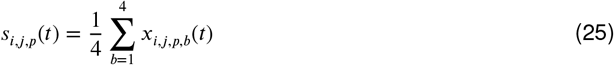

For each trial *i*, patch *p*, and time step *t*, we isolated the stochastic fluctuations around the generative mean by subtracting the mean generative luminance of that patch:

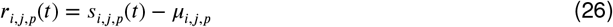

*r*_*i,j,p*_(*t*) reflects the momentary Gaussian luminance fluctuations. All kernels were computed using these residual signals.

Residual signals were aligned to the decision by defining the selected (*c*_*i,j*_) and non-selected (¬*c*_*i,j*_) patches:

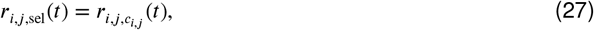

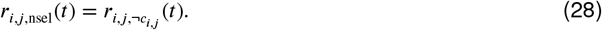

For the reverse correlation confidence analysis, we computed four traces:

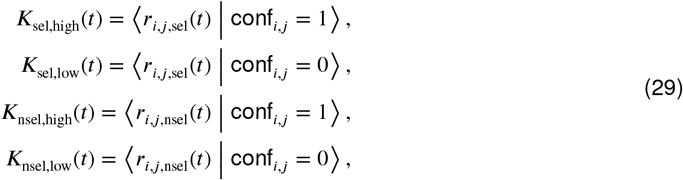

where ⟨⋅⟩ denotes averaging across trials. Each kernel therefore represents the average stochastic luminance fluctuation at time *t*, conditioned on whether the patch was selected or not and on the level of confidence.

To quantify the asymmetry in the contribution of luminance fluctuations to confidence, we fit a trial-level generalized linear mixed-effects regression model. The dependent variable, conf_*i,j*_, was a binary indicator of high confidence (1 = high). For each trial, we computed the average residual luminance fluctuations (i.e., frame-averaged luminance after subtraction of the generative mean) over the first 0.4 s for the selected and non-selected patches, yielding 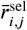 and 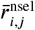. We also included the nominal luminance difference, Δ*L*_*i,j*_.

The model included fixed effects of 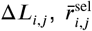, and 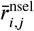, as well as a random intercept for subject (*b*_0*j*_):

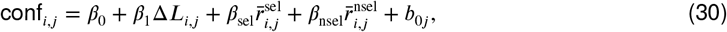

Models were fit separately for each task version (Choose-Brighter and Choose-Darker). To assess whether the contributions of the selected and non-selected patches were symmetric, we performed a linear hypothesis test of the null hypothesis *β*_sel_ + *β*_nsel_ = 0.

## Acknowledgments

A.Z. is grateful to Michael N. Shadlen for sharing resources and for many helpful discussions over the years, and to Gal Vishne for providing comments on an earlier version of the manuscript.

## Supplemental information

**Table S1.** Best-fitting parameter estimates for the hierarchical model.

| Subject | Task | $\kappa$ | $B$ | $T_{nd}$ | $\theta$ | $k_{urg}$ | $C_0$ | $k_{solv}$ |
| --- | --- | --- | --- | --- | --- | --- | --- | --- |
| S1 | Motion | 6.098 | 2.090 | 0.199 | 0.654 | 0.184 | 0.001 | 2.005 |
| S2 | Motion | 11.049 | 1.108 | 0.354 | 0.761 | 0.066 | -0.003 | 2.778 |
| S3 | Motion | 11.067 | 1.151 | 0.375 | 0.663 | 0.061 | 0.001 | 3.419 |
| S4 | Motion | 13.669 | 1.128 | 0.341 | 0.693 | 0.051 | 0.001 | 2.860 |
| S5 | Choose-Brighter | 22.085 | 1.540 | 0.352 | 0.949 | 0.028 | -0.000 | 2.001 |
| S6 | Choose-Brighter | 17.548 | 1.354 | 0.254 | 0.814 | 0.065 | -0.007 | 2.142 |
| S7 | Choose-Brighter | 27.035 | 1.060 | 0.320 | 0.845 | 0.009 | -0.005 | 2.619 |
| S8 | Choose-Brighter | 25.035 | 1.585 | 0.411 | 0.960 | 0.018 | 0.002 | 2.009 |
| S5 | Choose-Darker | 21.776 | 1.396 | 0.352 | 0.935 | 0.034 | -0.003 | 2.064 |
| S6 | Choose-Darker | 20.084 | 1.339 | 0.265 | 0.855 | 0.063 | 0.007 | 2.128 |
| S7 | Choose-Darker | 21.392 | 1.311 | 0.274 | 0.861 | 0.010 | -0.000 | 2.894 |
| S8 | Choose-Darker | 27.895 | 1.228 | 0.400 | 0.979 | 0.017 | 0.000 | 2.363 |

**Table S2.** Best-fitting parameter estimates for the integrated model.

| Subject | Task | $\theta$ | $\kappa$ | $B_h$ | $B_l$ | $T_{nd}$ | $p_{flip}$ | $C_0$ | $v_{max,l}$ | $v_{max,h}$ |
| --- | --- | --- | --- | --- | --- | --- | --- | --- | --- | --- |
| S1 | Motion | 0.837 | 5.704 | 2.326 | 5.289 | 0.141 | 0.098 | 0.026 | 4.362 | 7.985 |
| S2 | Motion | 1.121 | 7.041 | 1.690 | 2.652 | 0.195 | 0.041 | 0.006 | 5.065 | 5.857 |
| S3 | Motion | 0.725 | 7.799 | 2.015 | 3.214 | 0.184 | 0.016 | 0.020 | 5.028 | 6.700 |
| S4 | Motion | 1.335 | 10.826 | 1.280 | 2.657 | 0.317 | 0.024 | 0.002 | 6.241 | 4.521 |
| S5 | Choose-Brighter | 4.677 | 31.480 | 3.784 | 2.410 | 0.050 | 0.100 | 0.000 | 20.000 | 2.134 |
| S6 | Choose-Brighter | 9.038 | 1.974 | 3.682 | 3.612 | 0.381 | 0.038 | 0.014 | 7.147 | 7.716 |
| S7 | Choose-Brighter | 2.707 | 40.000 | 1.360 | 1.463 | 0.249 | 0.093 | -0.007 | 3.707 | 1.333 |
| S8 | Choose-Brighter | 5.166 | 36.382 | 4.527 | 2.492 | 0.050 | 0.100 | 0.001 | 20.000 | 1.799 |
| S5 | Choose-Darker | 3.936 | 25.502 | 3.451 | 2.350 | 0.078 | 0.100 | 0.000 | 20.000 | 2.679 |
| S6 | Choose-Darker | 2.425 | 14.962 | 1.445 | 1.680 | 0.211 | 0.023 | 0.008 | 10.805 | 3.086 |
| S7 | Choose-Darker | 2.851 | 40.000 | 1.984 | 1.959 | 0.089 | 0.100 | -0.004 | 3.168 | 1.334 |
| S8 | Choose-Darker | 4.770 | 33.046 | 3.740 | 2.145 | 0.081 | 0.083 | 0.000 | 20.000 | 2.247 |

**Figure S1.**
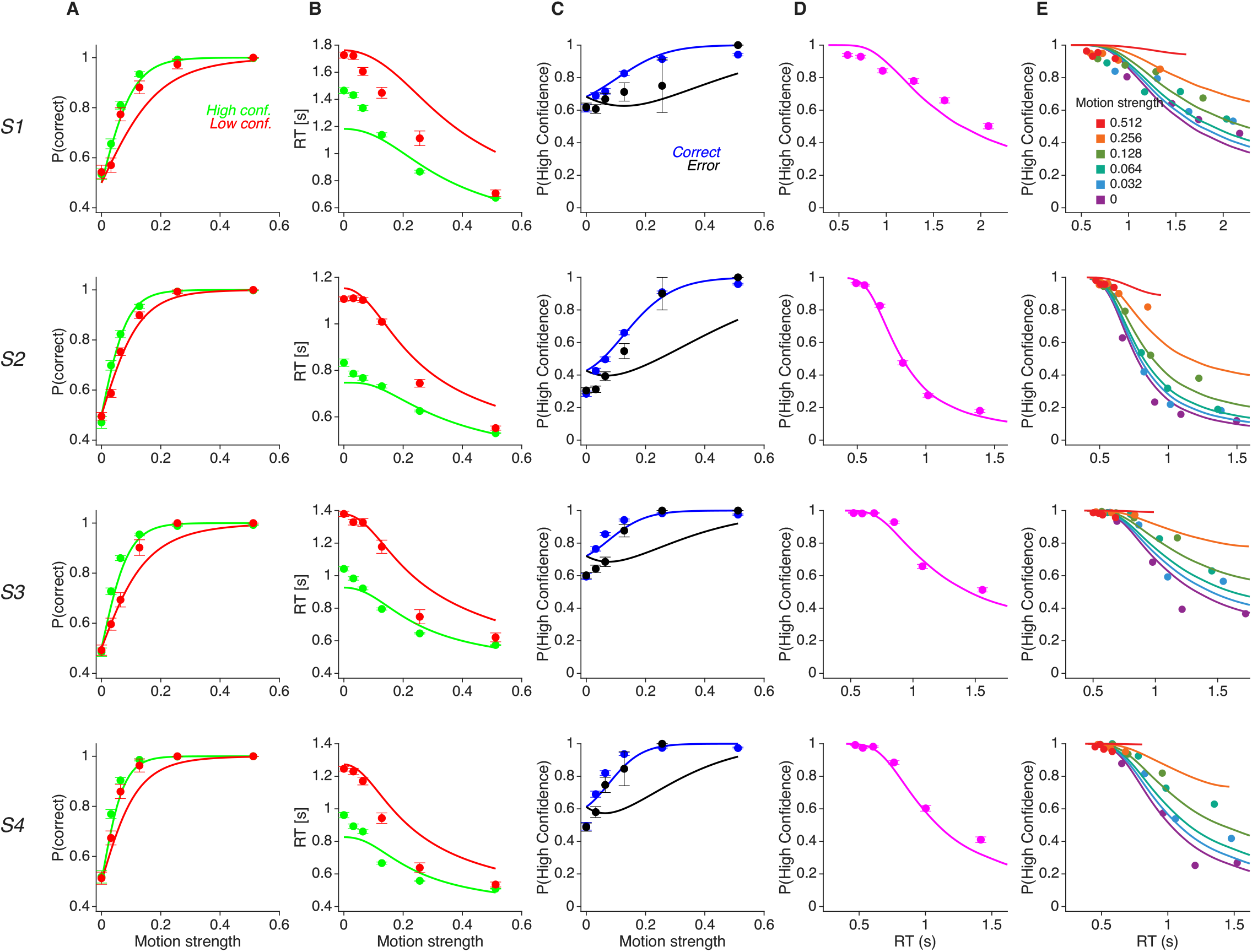
Fits of the hierarchical model with *rho* = −0.7071 to the motion task data. Same conventions as in Fig. 5.

**Figure S2.**
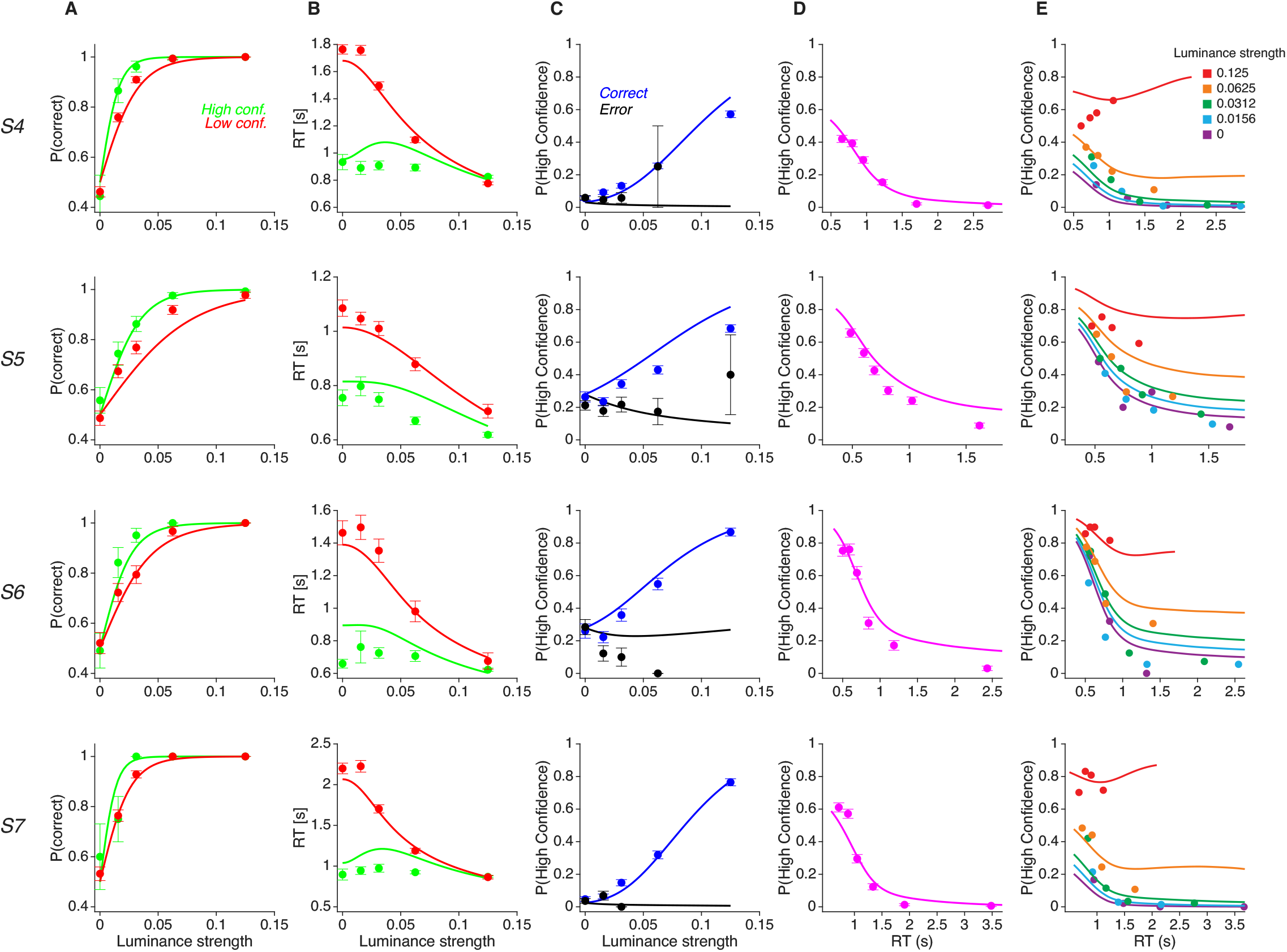
Behavior in the luminance (Choose-Darker) task and fits of the hierarchical model. Same conventions as in Fig. 5.

**Figure S3.**
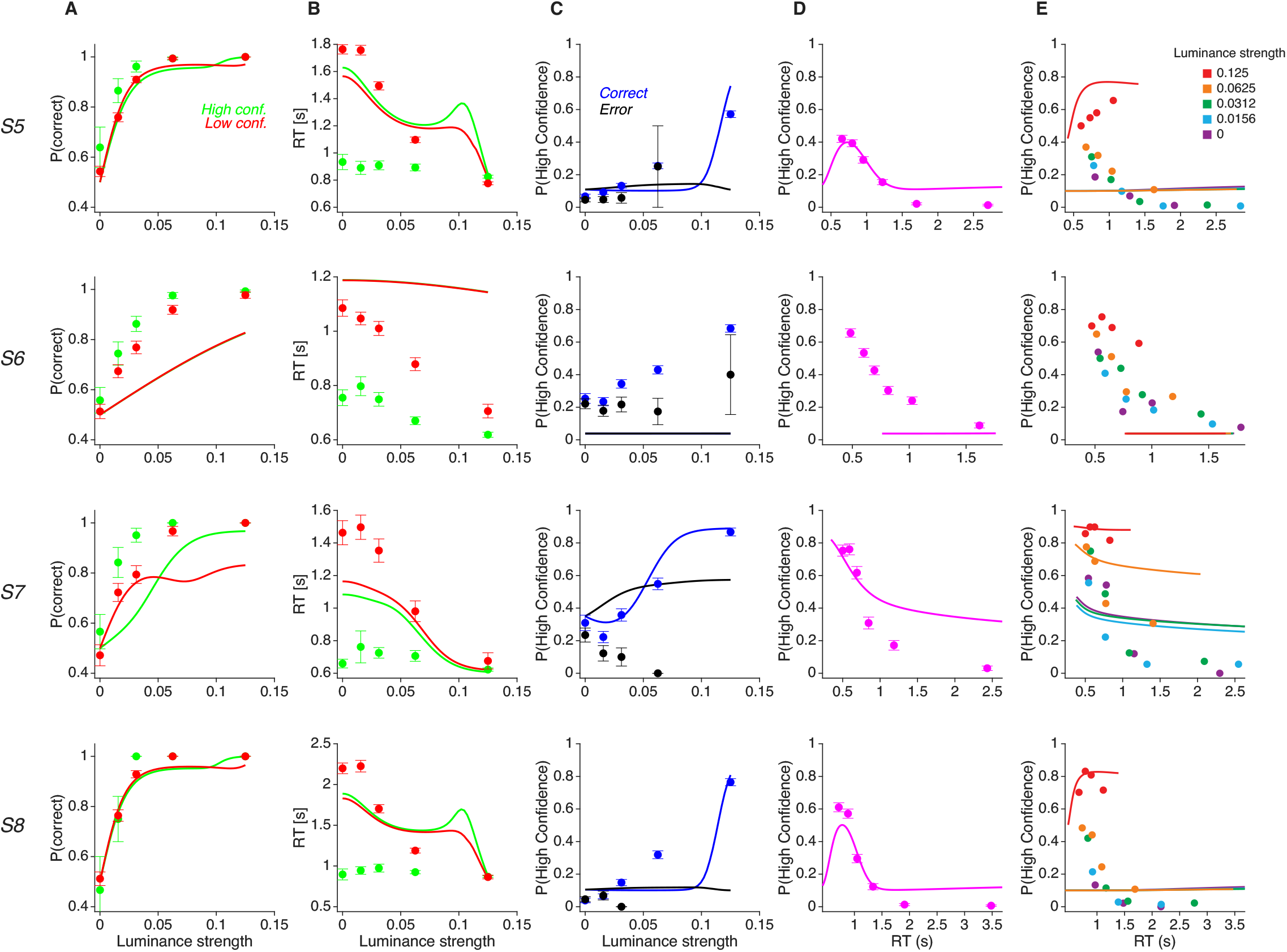
Behavior in the luminance (Choose-Darker) task and fits of the flat competition model. Same conventions as in Fig. 5.

**Figure S4.**
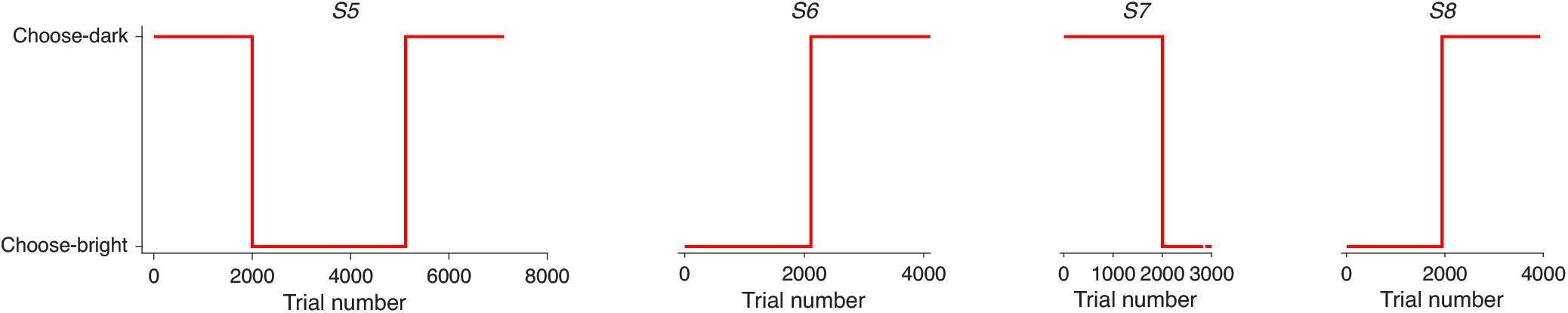
Trials per participant and task version in the luminance task. Participants completed two versions of the luminance task (Choose-Brighter and Choose-Darker) in counterbalanced order. Each participant first completed 2,000 trials of one version (across two sessions) before switching to the other version. Each panel shows the task version performed by each participant (S4–S7). Participant S4 returned to the Choose-Darker version after completing the Choose-Brighter version.

**Figure S5.**
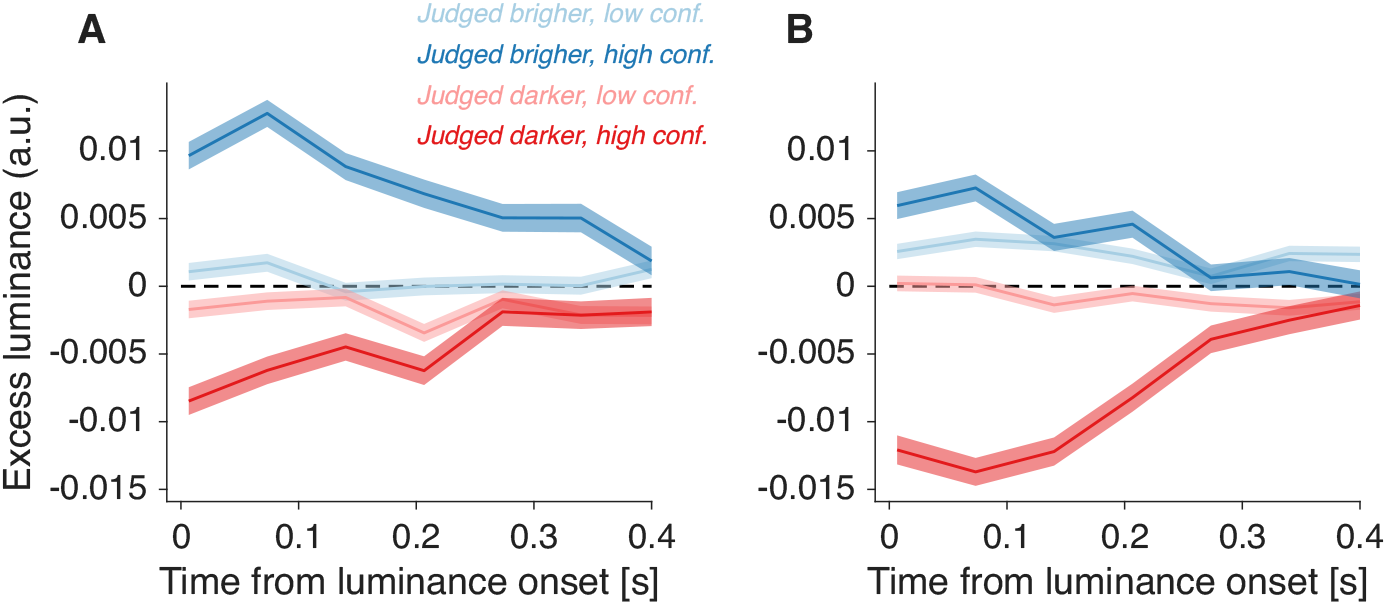
Psychophysical kernels of luminance fluctuations including all trials. Same analysis and conventions as in Fig. 10, but including all trials regardless of whether participants had previously completed the other version of the task. Results are largely similar to those in Fig. 10, although the separation between high- and low-confidence traces is larger for the non-selected patch (red in (A), blue in (B)). This may reflect learning effects or transfer between task versions (e.g., prior experience with the Choose-Brighter task influencing performance in the other version). Crucially, the main result remains: fluctuations in the selected patch are more informative about confidence than those in the non-selected patch.

## References

Acerbi L, Ma WJ. Practical Bayesian optimization for model fitting with Bayesian adaptive direct search. arXiv preprint arXiv:170504405. 2017;.

Ais J, Zylberberg A, Barttfeld P, Sigman M. Individual consistency in the accuracy and distribution of confidence judgments. Cognition. 2016; 146:377–386.

Audley R. A stochastic model for individual choice behavior. Psychological review. 1960; 67(1):1.

Baird B, Cieslak M, Smallwood J, Grafton ST, Schooler JW. Regional white matter variation associated with domain-specific metacognitive accuracy. Journal of Cognitive Neuroscience. 2015; 27(3):440–452.

Barttfeld P, Wicker B, McAleer P, Belin P, Cojan Y, Graziano M, Leiguarda R, Sigman M. Distinct patterns of functional brain connectivity correlate with objective performance and subjective beliefs. Proceedings of the National Academy of Sciences. 2013; 110(28):11577–11582.

Van den Berg R, Zylberberg A, Kiani R, Shadlen MN, Wolpert DM. Confidence is the bridge between multi-stage decisions. Current Biology. 2016; 26(23):3157–3168.

Bona S, Silvanto J. Accuracy and confidence of visual short-term memory do not go hand-in-hand: Behavioral and neural dissociations. PLoS One. 2014; 9(3):e90808.

Brus J, Aebersold H, Grueschow M, Polania R. Sources of confidence in value-based choice. Nature communications. 2021; 12(1):7337.

Chen H, Heathcote A, Osth AF, Chen H. Linear Ballistic Accumulator Models of Confidence and Response Time. 2026;.

Churchland AK, Kiani R, Shadlen MN. Decision-making with multiple alternatives. Nature neuroscience. 2008; 11(6):693–702.

David AS, Bedford N, Wiffen B, Gilleen J. Failures of metacognition and lack of insight in neuropsychiatric disorders. Nature Reviews Neurology. 2012; 8(8):412–426.

Del Cul A, Dehaene S, Reyes P, Bravo E, Slachevsky A. Causal role of prefrontal cortex in the threshold for access to consciousness. Brain. 2009; 132(9):2531–2540.

van Den Berg R, Anandalingam K, Zylberberg A, Kiani R, Shadlen MN, Wolpert DM. A common mechanism underlies changes of mind about decisions and confidence. Elife. 2016; 5:e12192.

DePasquale B, Brody CD, Pillow JW, Neural population dynamics underlying evidence accumulation in multiple rat brain regions. Neuroscience; 2021. http://biorxiv.org/lookup/doi/10.1101/2021.10.28.465122, doi: 10.1101/2021.10.28.465122.

Dou W, Afrakhteh S, Samaha J. Metacognitive Introspection Alters the Dynamics of Pre-Decisional Neural Evidence Accumulation. BioRxiv. 2024; p. 2024–11.

Double KS, Birney DP. Confidence judgments interfere with perceptual decision making. Scientific Reports. 2024; 14(1):14133.

Double KS, Pinkus RT, Landfear R, Carnegie P, Dowell S, Goldwater MB. Rating our certainty: how confidence judgments amplify belief polarization. Psychological Research. 2026; 90(1):25.

Fleming SM, Daw ND. Self-evaluation of decision-making: A general Bayesian framework for metacognitive computation. Psychological review. 2017; 124(1):91.

Fleming SM, Ryu Ji, Golfinos JG, Blackmon KE. Domain-specific impairment in metacognitive accuracy following anterior prefrontal lesions. Brain. 2014; 137(10):2811–2822.

Fleming SM, Weil RS, Nagy Z, Dolan RJ, Rees G. Relating introspective accuracy to individual differences in brain structure. Science. 2010; 329(5998):1541–1543.

Gold JI, Shadlen MN. The neural basis of decision making. Annual review of neuroscience. 2007; 30.

Graziano M, Sigman M. The spatial and temporal construction of confidence in the visual scene. PLoS One. 2009; 4(3):e4909.

Hanks TD, Kopec CD, Brunton BW, Duan CA, Erlich JC, Brody CD. Distinct relationships of parietal and prefrontal cortices to evidence accumulation. Nature. 2015; 520(7546):220–223.

Hellmann S, Zehetleitner M, Rausch M. Simultaneous modeling of choice, confidence, and response time in visual perception. Psychological review. 2023; 130(6):1521.

Howard IS, Ingram JN, Wolpert DM. A modular planar robotic manipulandum with end-point torque control. Journal of neuroscience methods. 2009; 181(2):199–211.

Hyafil A, Moreno-Bote R. Breaking down hierarchies of decision-making in primates. Elife. 2017; 6:e16650.

Kiani R, Corthell L, Shadlen MN. Choice certainty is informed by both evidence and decision time. Neuron. 2014; 84(6):1329–1342.

Komura Y, Nikkuni A, Hirashima N, Uetake T, Miyamoto A. Responses of pulvinar macaque neurons reflect a subject’s confidence in visual categorization. Nature Neuroscience. 2013; 16(6):749–755.

Lak A, Costa RM, Romberg C, Koulakov AA, Mainen ZF, Kepecs A. Orbitofrontal cortex is required for optimal waiting based on decision confidence. Neuron. 2014; 84(1):190–201.

Latimer KW, Yates JL, Meister ML, Huk AC, Pillow JW. Single-trial spike trains in parietal cortex reveal discrete steps during decision-making. Science. 2015; 349(6244):184–187.

Lau HC, Passingham RE. Relative blindsight in normal observers and the neural correlate of visual consciousness. Proceedings of the National Academy of Sciences. 2006; 103(49):18763–18768.

Litwin P, Paulewicz B, Siedlecka M. Reporting confidence decreases response and change-of-mind accuracy in a perceptual decision task. Journal of Experimental Psychology: Human Perception and Performance. 2025;.

Lorteije JA, Zylberberg A, Ouellette BG, De Zeeuw CI, Sigman M, Roelfsema PR. The formation of hierarchical decisions in the visual cortex. Neuron. 2015; 87(6):1344–1356.

Mazor M, Maimon-Mor RO, Charles L, Fleming SM. Paradoxical evidence weighting in confidence judgments for detection and discrimination. Attention, Perception, & Psychophysics. 2023; 85(7):2356–2385.

Meyniel F, Sigman M, Mainen Z. Confidence as Bayesian Probability: From Neural Origins to Behavior. Neuron. 2015 Oct; 88(1):78–92. https://linkinghub.elsevier.com/retrieve/pii/S0896627315008284, doi: 10.1016/j.neuron.2015.09.039.

Palmer EC, David AS, Fleming SM. Effects of age on metacognitive efficiency. Consciousness and Cognition. 2014; 28:151–160.

Paz L, Insabato A, Zylberberg A, Deco G, Sigman M. Confidence through consensus: a neural mechanism for uncertainty monitoring. Scientific Reports. 2016 Feb; 6(1):21830. https://www.nature.com/articles/srep21830, doi: 10.1038/srep21830.

Peters MA, Thesen T, Ko YD, Maniscalco B, Carlson C, Davidson M, Doyle W, Kuzniecky R, Devinsky O, Halgren E, et al. Perceptual confidence neglects decision-incongruent evidence in the brain. Nature human behaviour. 2017; 1(7):0139.

Petrusic WM, Baranski JV. Judging confidence influences decision processing in comparative judgments. Psychonomic bulletin & review. 2003; 10(1):177–183.

Pouget A, Drugowitsch J, Kepecs A. Confidence and certainty: distinct probabilistic quantities for different goals. Nature neuroscience. 2016; 19(3):366–374.

Rahnev D. Confidence-accuracy dissociations in perceptual decision making. Available at SSRN 5828455. 2026;.

Rahnev D, Denison RN. Suboptimality in perceptual decision making. Behavioral and brain sciences. 2018; 41:e223.

Rahnev D, Maniscalco B, Graves T, Huang E, de Lange FP, Lau H. Attention induces conservative subjective biases in visual perception. Nature Neuroscience. 2011; 14(12):1513–1515.

Ratcliff R, McKoon G. The diffusion decision model: theory and data for two-choice decision tasks. Neural computation. 2008; 20(4):873–922.

Ratcliff R, Starns JJ. Modeling confidence and response time in recognition memory. Psychological review. 2009; 116(1):59.

Ratcliff R, Starns JJ. Modeling confidence judgments, response times, and multiple choices in decision making: recognition memory and motion discrimination. Psychological review. 2013; 120(3):697.

Roitman JD, Shadlen MN. Response of neurons in the lateral intraparietal area during a combined visual discrimination reaction time task. Journal of neuroscience. 2002; 22(21):9475–9489.

Rounis E, Maniscalco B, Rothwell JC, Passingham RE, Lau H. Theta-burst transcranial magnetic stimulation to the prefrontal cortex impairs metacognitive visual awareness. Cognitive Neuroscience. 2010; 1(3):165–175.

Sepulveda P, Usher M, Davies N, Benson AA, Ortoleva P, De Martino B. Visual attention modulates the integration of goal-relevant evidence and not value. Elife. 2020; 9:e60705.

Shadlen MN, Kiani R, Hanks TD, Churchland AK. An intentional framework. Better than conscious. 2008; p. 71–101.

Shan H, Moreno-Bote R, Drugowitsch J. Family of closed-form solutions for two-dimensional correlated diffusion processes. Physical Review E. 2019; 100(3):032132.

Song C, Kanai R, Fleming SM, Weil RS, Schwarzkopf DS, Rees G. Relating inter-individual differences in metacognitive performance on different perceptual tasks. Consciousness and Cognition. 2011; 20(4):1787–1792.

Steinemann NA, Stine GM, Trautmann EM, Zylberberg A, Wolpert DM, Shadlen MN. Direct observation of the neural computations underlying a single decision. bioRxiv. 2022; p. 2022–05.

Toso A, Arazi A, de la Rocha J, Tsetsos K, Donner TH. Competing Neural Decision Variables in Human Frontal Cortex Shape Decision Confidence. bioRxiv. 2025; p. 2025–10.

Vickers D. Decision processes in visual perception. Academic Press; 1979.

Vivar Lazo M. Cortical Population Dynamics Underlying Choice, Reaction Time, and Confidence. PhD thesis, Johns Hopkins University; 2024.

Vivar-Lazo M, Fetsch CR. Neural basis of concurrent deliberation toward a choice and confidence judgment. Nature neuroscience. 2025; p. 1–12.

Vlassova A, Donkin C, Pearson J. Unconscious information changes decision accuracy but not confidence. Proceedings of the National Academy of Sciences. 2014; 111(45):16214–16218.

Wilimzig C, Tsuchiya N, Fahle M, Einhäuser W, Koch C. Spatial attention increases performance but not subjective confidence in a discrimination task. Journal of Vision. 2008; 8(5):7–7.

Yeung N, Summerfield C. Metacognition in human decision-making: confidence and error monitoring. Philosophical Transactions of the Royal Society B: Biological Sciences. 2012; 367(1594):1310–1321.

Zylberberg A, Barttfeld P, Sigman M. The construction of confidence in a perceptual decision. Frontiers in integrative neuroscience. 2012; 6:79.

Zylberberg A, Lorteije JA, Ouellette BG, De Zeeuw CI, Sigman M, Roelfsema P. Serial, parallel and hierarchical decision making in primates. Elife. 2017; 6:e17331.

Zylberberg A, Shadlen MN. Cause for pause before leaping to conclusions about stepping. BioRxiv. 2016; p. 085886.

Zylberberg A, Shadlen MN. A population representation of the confidence in a decision in the parietal cortex. Cell reports. 2025; 44(4).

